# Kinetic fingerprints differentiate anti-Aβ therapies

**DOI:** 10.1101/815308

**Authors:** Sara Linse, Tom Scheidt, Katja Bernfur, Michele Vendruscolo, Christopher M. Dobson, Samuel I. A. Cohen, Eimantas Sileikis, Martin Lundquist, Fang Qian, Tiernan O’Malley, Thierry Bussiere, Paul H. Weinreb, Catherine K. Xu, Georg Meisl, Sean R. A. Devenish, Tuomas P. J. Knowles, Oskar Hansson

## Abstract

The amyloid cascade hypothesis, according to which the self-assembly of amyloid-β peptide (Aβ) is a causative process in Alzheimer’s disease, has driven many therapeutic efforts for the past 20 years. Failures of clinical trials investigating Aβ-targeted therapies have been interpreted as evidence against this hypothesis, irrespective of the characteristics and mechanisms of action of the therapeutic agents, which are highly challenging to assess. We bring together kinetic analysis with quantitative binding measurements to address the mechanisms of action of four clinical stage anti-Aβ antibodies, aducanumab, gantenerumab, bapineuzumab and solanezumab. We reveal and quantify the striking differences of these antibodies on the aggregation kinetics and on the production of oligomeric aggregates, and link these effects to the affinity and stoichiometry of each antibody for monomeric and fibrillar forms of Aβ. Our results uncover that, uniquely amongst these four antibodies, aducanumab dramatically reduces the flux of oligomeric forms of Aβ.

---

Alzheimer’s disease affects nearly 50 million people worldwide with an overall cost corresponding to more than 1% of the global economy. The amyloid hypothesis (1,2) has motivated the development of many therapeutic antibodies targeting different species of the Aβ peptide, with most of these immunotherapeutic approaches focusing on achieving a global reduction in the concentration of Aβ aggregates or a corresponding delay in the overall Aβ aggregation rate (3). Increasing evidence, however, indicates that different Aβ aggregates may have different pathological effects (4,5). In particular, biophysical, cellular and histological data have highlighted a more pronounced toxicity of oligomeric intermediates of lower molecular weight, relative to that of large fibrillar species. It is therefore critical to determine the effects of potential anti-Aβ therapeutics on the underlying microscopic steps of Aβ aggregation (5,6), since many overall interventions are not likely to affect the levels of the most toxic species generated through this process, and might consequently be less clinically effective.

Specifically, studies of the Aβ aggregation mechanism have revealed a multistep process that includes primary nucleation of monomers into aggregates, growth of aggregates by monomer addition and autocatalytic aggregate multiplication by secondary nucleation of monomers at the surfaces of existing fibrils (7). Above a critical concentration of fibrils, this latter process becomes the dominant source of new low molecular weight Aβ oligomers. Secondary nucleation thereby connects the production of oligomers to the concentrations of both monomers and fibrils through a non-linear feedback network. There are two fundamental approaches to reduce the flux of oligomeric species: first by directly inhibiting the molecular reaction through which they form, and second by removal of the reactants, including the catalytic fibril surfaces. We present here a strategy based on a quantitative molecular analysis to assess the effects of four clinical stage antibodies, aducanumab (8), gantenerumab (9,10), bapineuzumab (11) and solanezumab (12), as murine analogs ^ch^aducanumab, ^ch^gantenerumab, 3D6 and m266, respectively, on both of these potential therapeutic modes of action. Clinical trials of Bapineuzumab and Solanezumab have been discontinued, while those of aducanumab and gantenerumab are still active.

First, we assessed the potency of the antibodies on the direct inhibition of secondary nucleation using chemical kinetics (5,7,13). The action of an inhibitor is assessed through the global analysis of aggregation reaction time courses (5,6, Fig. S1), which are characterized by a lag-phase, a so-called growth phase and a final plateau. Crucially, all three underlying microscopic processes occur during all three phases, albeit at different rates, as governed by the rate constants and concentrations of reacting species at each point in time (14,15), implying that simple observations of the overall aggregation behavior do not readily allow the changes in molecular mechanisms to be determined. In particular, both elongation and secondary nucleation exhibit their maximum rates during the macroscopic growth phase; the total nucleation rate is highest close to the mid-point (t_0.5_) of the reaction where both fibrils and monomers are present at approximately equal concentrations.

The aggregation of the 42-residue form of Aβ (Aβ42) was initiated by a temperature jump from 0 to 37°C, thus creating supersaturated solutions of Aβ42 monomers in the presence or absence of antibodies in pure buffer (Fig. 1, S2-S5) or in human cerebrospinal fluid (CSF, Fig. S6, 16,17). For each dataset, recorded at constant Aβ42 concentration and increasing concentrations of each antibody, we performed a fit to the integrated rate law describing the Aβ42 aggregation process where one of the molecular rate constants, kn, k2 or k+ for primary nucleation, secondary nucleation and elongation, respectively, was allowed to vary upon addition of the antibody (16, Fig. 1). Moreover, experiments were conducted in the presence of fixed low (Fig. S7,S8) or high (Fig. 1, first column) concentrations of pre-formed seed-aggregates in the initial reactant solution, thereby bypassing primary or both primary and secondary nucleation to probe specifically the effects on secondary nucleation and/or elongation. All the experiments were run in a blinded manner using the four anti-Aβ antibodies as well as an isotype control antibody P1.17.

**Figure 1.**
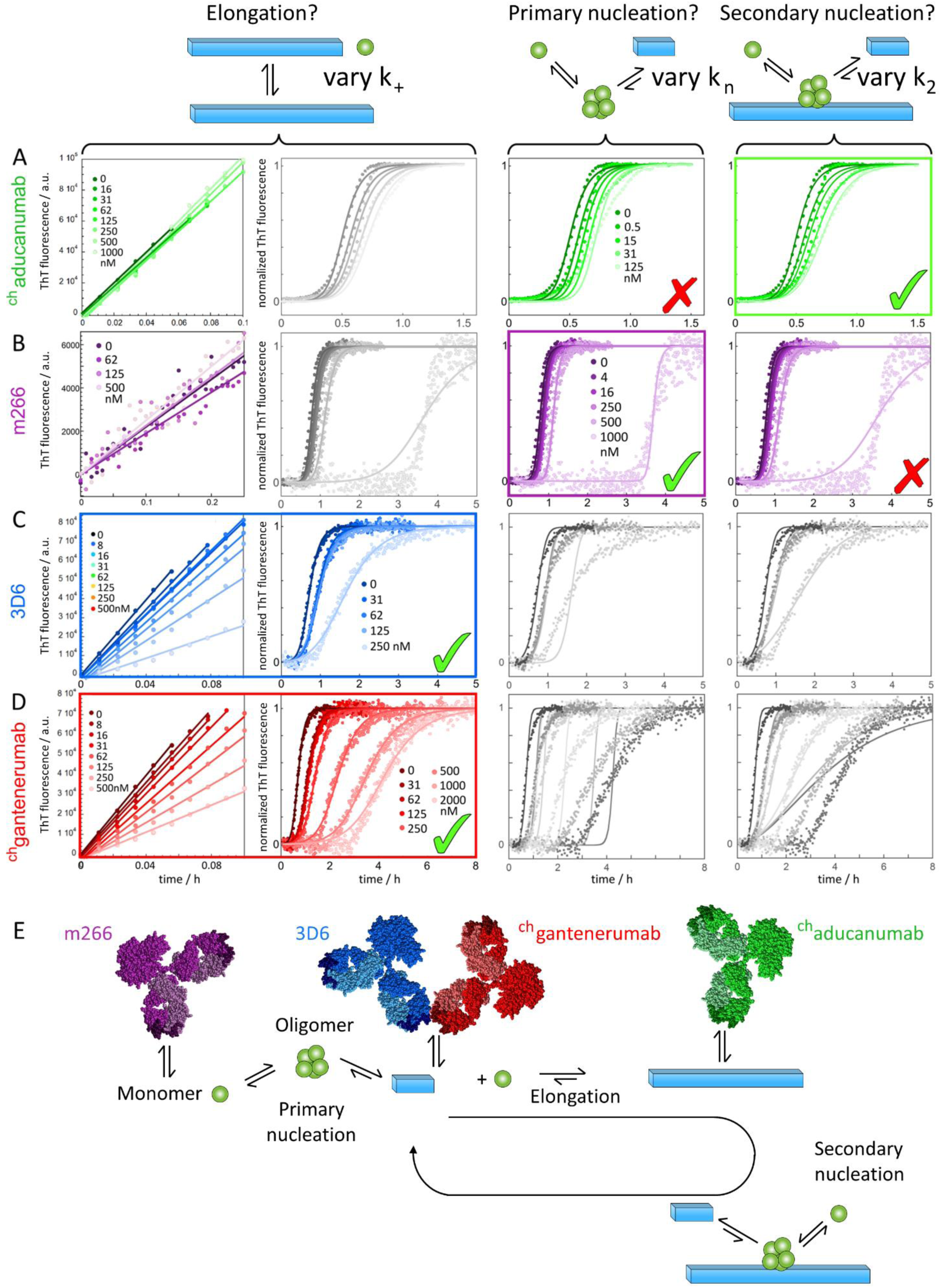
Effects of the antibodies on the kinetics of aggregation of Aβ42 in the presence and absence of pre-formed fibrils. **(A-D)** Thioflavin T (ThT) fluorescence as a function of time for reactions starting from 3-4 µM recombinant Aβ1-42 in 20 mM HEPES/NaOH, 140 mM NaCl, 1 mM CaCl_2_, pH 8.0 in the absence and presence of ^ch^aducanumab (**A**), m266 (**B**), 3D6 (**C**) or ^ch^gantenerumab (**D**). The first column shows data obtained for Aβ1-42 monomer in the presence of 30% pre-formed Aβ1-42 fibrils. The last three columns show data obtained for monomeric solutions in the absence of pre-formed fibrils. The colour codes for the antibody concentrations in nM are given on each row. The data are shown in normalized form, with the non-normalized data in Fig. S3. Data in the presence of isotype control are shown in Fig. S3C. The solid lines are fits to the data and assume in the left two columns global values for k_2_ and k_n_ and curve-specific values for k_+_, in the third column global values for k_+_ and k_n_ and curve-specific values for k_2,_ and in the right column global values for k_2_ and k_+_ and curve-specific values for k_2_. The best fit in each case is indicated by a green tick. Grey colour indicates discarded mechanisms based on the results of heavy seeded data (first column). Note that the x-axes cover different ranges depending on the magnitude of the effect of each antibody. Data and analysis for additional concentrations of each antibody are shown in Fig. S3 and S4. **(E)** Schematic illustration of the different microscopic steps in the aggregation that are primarily affected by the four antibodies. Structural models were prepared using MOLMOL (26).

We found that ^ch^aducanumab selectively and dramatically reduces the secondary nucleation rate of Aβ42 (Fig. 1A), lowering the effective rate constant for this process by ca. 40% even at the lowest concentrations of antibody tested (250 pM, Fig. S4). By contrast, the data show that m266 selectively inhibits primary nucleation (Fig. 1C). For both of these antibodies, no changes are detected to the highly pre-seeded aggregation reaction when high concentrations of antibody are introduced into the reactions, highlighting that the growth of pre-existing aggregates is largely unchanged (Fig. 1). Moreover, 3D6 and ^ch^gantenerumab both act predominantly by reducing the growth of the fibrillar aggregates under both pre-seeded and unseeded reaction conditions (Fig. 1B, 1D). For comparison, the isotype control antibody has no detectable effect even at high concentration (Fig. S3). In order to extend these findings to physiological conditions, we used CSF in the aggregation assay. In this case, Aβ42 displayed an extended lag phase consistent with earlier findings (Fig. S6, ref. 17,18). However, each antibody was found to predominantly inhibit the same step as in buffer (Fig. 1, S6,S7), with variations in the values of the rate constants reflecting the change of environment from buffer to CSF.

We next verified the reduction in oligomer production predicted to originate from inhibition of secondary nucleation by performing direct measurements of the oligomer concentrations. To this effect, we quantified the concentration of oligomers present at the half time of the aggregation reaction using MALDI mass spectrometry and size-exclusion chromatography (SEC), after the addition of a known amount of isotope standard (^15^N-Aβ42) and proteolytic digestion (19; Fig. 2). The results reveal that ^ch^aducanumab causes a clear reduction of the free oligomer concentration, in direct agreement with the observed reduction in secondary nucleation, while a smaller reduction is observed with 3D6 and in particular with ^ch^gantenerumab, neither of which significantly inhibits secondary nucleation. Interestingly, a fraction of Aβ42 elutes together with ^ch^aducanumab in the void; given its low affinity of ^ch^aducanumab for Aβ42 monomers this most likely represents bound oligomers (Fig. S9). Such direct binding of oligomers could potentially prevent their further conversion to growth-competent fibrils, but could in turn increase their population, although the bound oligomers may be de-toxified by such capture. For m266 we observe a very large amount of Aβ42 eluting in early size exclusion chromatography fractions (Fig. S10), consistent with the observation that this antibody binds with high affinity to the monomer (24).

**Figure 2.**
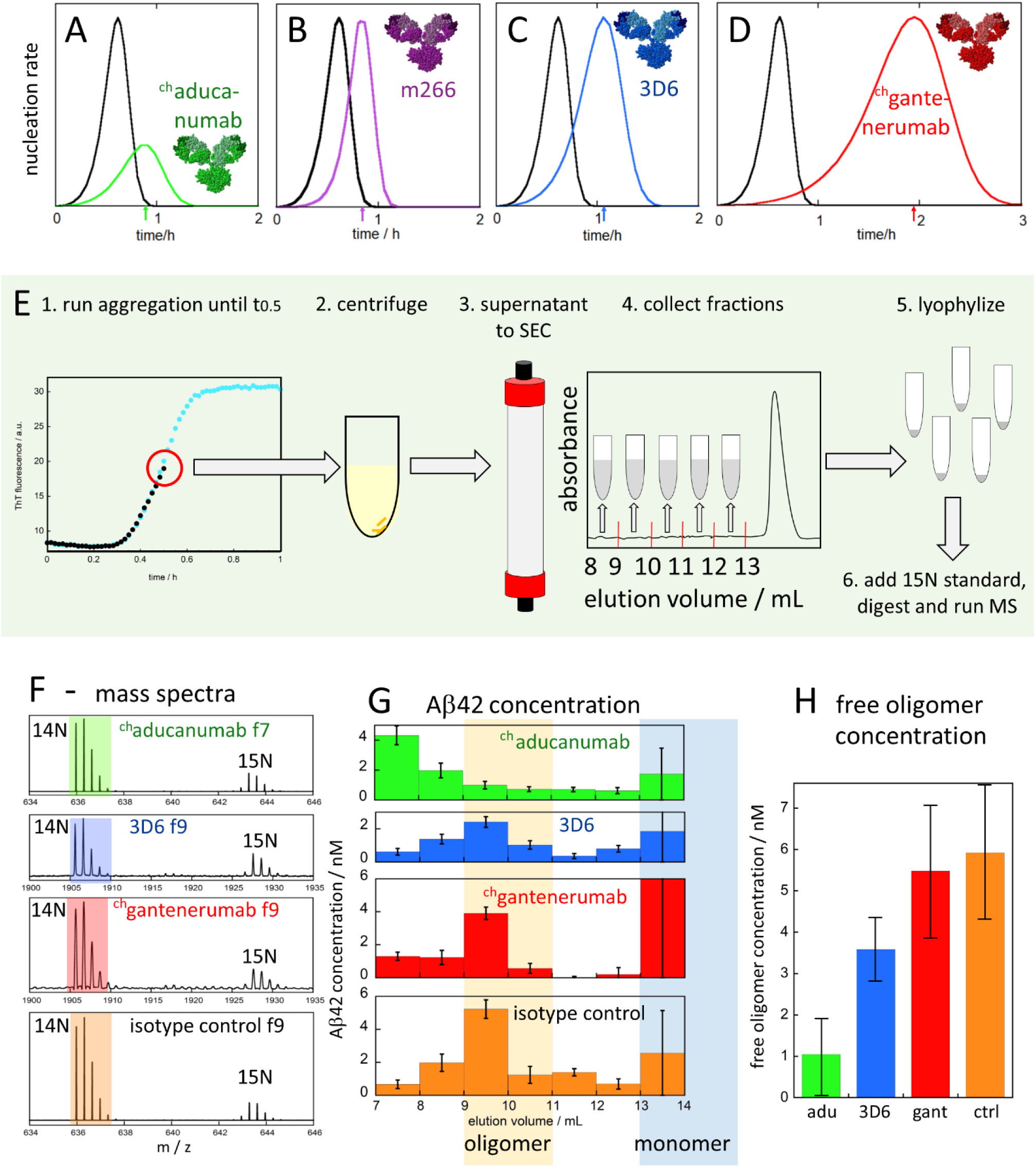
Effects of the antibodies on the production of Aβ42 oligomer. (**A-D**) Predicted effects of the four antibodies on the Aβ1-42 nucleation rate based on fitted rate constants. The arrows on the time axis point to t_half_ of the macroscopic aggregation curve. (**E**) Outline of the oligomer quantification experiment. Samples were collected at t_half_, the point in time where the ThT fluorescence was half-way in between the baseline and plateau values for each respective reaction. The samples were centrifuged, the supernatant separated by SEC and 1 mL fractions collected from the void to the start of the monomer peak, lyophilized, supplemented with 1 nM ^15^N-Aβ(M1-42) as an isotope standard, digested and analysed by mass spectrometry. (**F**) Examples of data showing the DSGYEVHHQKLVFFAE peptide pair where the ^14^N peptide has a monoisotopic mass of 1905.91 Da and the ^15^N peptide a mass of 1927.91 Da. These two peptides are chemically equivalent and can be used for quantification. Example data for one fraction (f) from reactions in the presence of^ch^aducanumab (f7), 3D6 (f9), ^ch^gantenerumab (f9) and isotype control (f9). (**G**) Observed size distribution of oligomers from one experiment. (**H**) Free oligomer concentration eluting from samples collected at t_half_ in the presence of each antibody. The data shown are averages of fraction 9+10 from two repeats and the error bars include the values in both repeats plus estimated maximum fraction collection errors. The data for m266 are not included in the graph since in this case a very large amount of Aβ1-42 elutes together with the antibody in the column void (Fig. S10).

The specific inhibition of secondary nucleation (Fig. 1), and the corresponding reduction in free oligomer concentration (Fig. 2), identified in the presence of ^ch^aducanumab implies that this antibody’s activity is likely to be driven predominantly by interactions with species unique to secondary nucleation, i.e. the fibril surface, rather than interactions with soluble species which are involved in both primary and secondary nucleation. To verify the molecular species on which this antibody acts, therefore, we performed experiments in which newly formed Aβ42 fibrils, produced in the presence or absence of ^ch^aducanumab, were diluted and added to freshly prepared monomer solutions (Fig. S11). We observed that pristine fibrils that had not been exposed to ^ch^aducanumab at any stage enhance the process of secondary nucleation and increase the rate of aggregation, whereas pre-formed fibrils that had been generated in the presence of 0.1 molar equivalents of ^ch^aducanumab accelerate aggregation to a significantly smaller extent. Kinetic analysis (Fig. S11) reveals a reduction of approximately ca. 33% in the apparent rate constant for secondary nucleation in this latter case, a value that is similar to that identified from the analysis in Fig. 1 (see also Fig. S4), showing that a significant fraction of the inhibitory effect on secondary nucleation originates from interaction of the antibody with fibrillar aggregates.

In order to further investigate the origin of the kinetic inhibition and to assess additional modes of action through reduction in the concentrations of the reactants for secondary nucleation, we probed directly the stoichiometry and affinity of the interactions between each antibody and Aβ42 monomers and fibrils. These interactions were investigated by monitoring changes in the diffusion coefficients of the equilibrated species through microfluidic diffusional sizing (20,21; Fig. 3). The data reveal that ^ch^aducanumab has a low affinity for Aβ42 monomers (pK_D_ = -^10^logK_D_ = ^10^logK = 5.04±0.08, K_D_=9±2 µM, the error given is the standard deviation), but a very high affinity for fibrils (pK_D_ = 9.0±0.1, K_D_=1.0±0.2 nM). The affinity we obtain here in solution is a factor of 5 lower than for surface-immobilized fibrils (24). These findings imply a remarkable specificity ratio of over four orders of magnitude for fibril binding over monomer binding, potentially a result of avidity effects since the epitope, residues 3-7, (24) is exposed in monomers as well as fibrils; this segment is relatively disordered and not included in the ssNMR structure of Aβ42 fibrils (22; Fig. 3B). In addition, the diffusion data provide information on the stoichiometry of binding. We find one ^ch^aducanumab molecule per ca. 5 Aβ42 monomers in the fibril (Fig 3C, S13). Interestingly, the width of each antigen-binding region of an IgG is indeed similar to the summed pitch of 5 planes of an Aβ42 fibril (Fig. 3B), and the measured value of the stoichiometry implies that fibrils can become fully coated with ^ch^aducanumab along their entire length, which would effectively interfere with secondary nucleation at the fibril surface. The remarkable specificity of ^ch^aducanumab in suppressing secondary nucleation with no observable effect on elongation (Fig. 1) redirects a significant part of the reactive flux from secondary nucleation to elongation.

**Figure 3.**
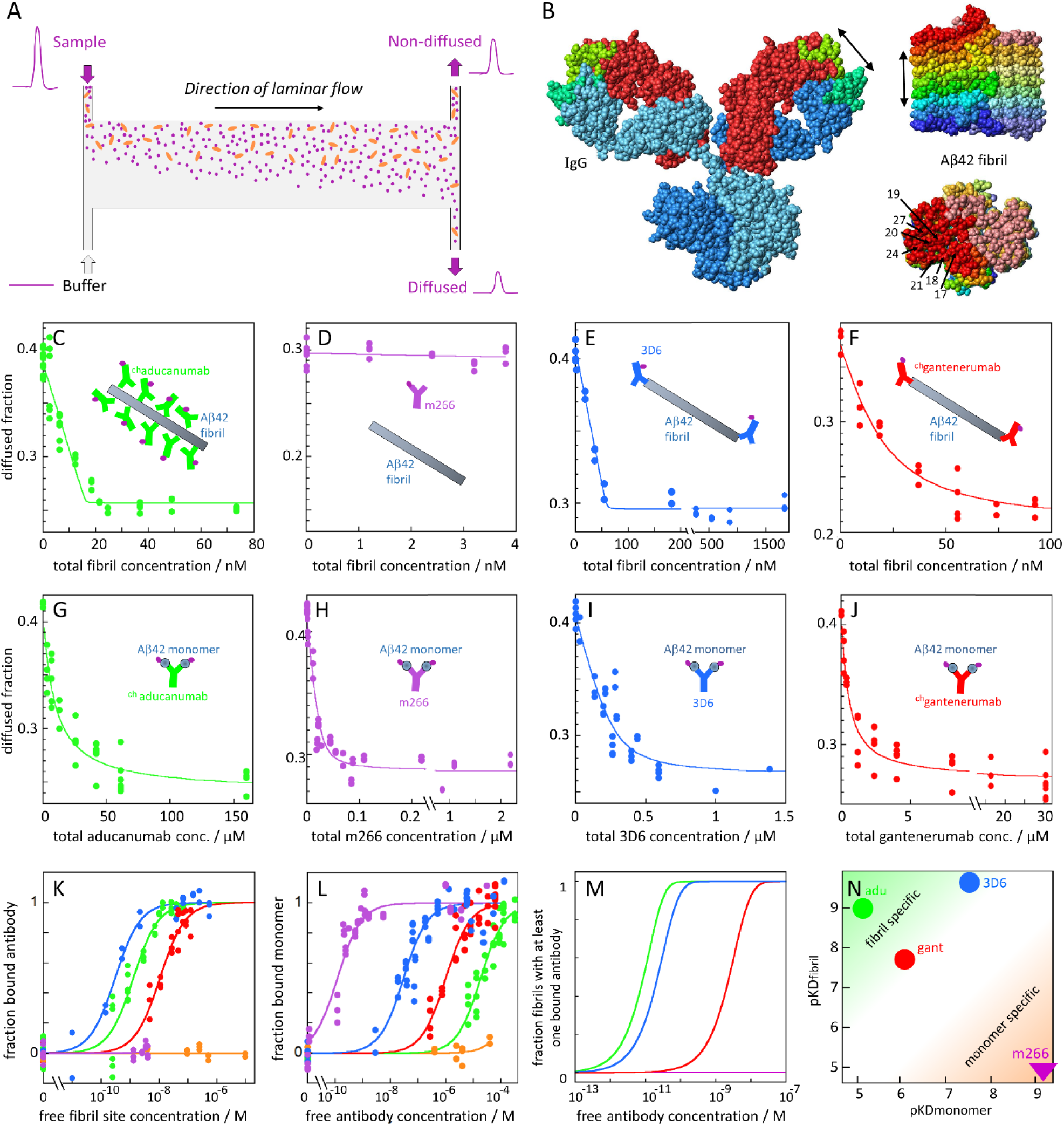
Binding affinity of the antibodies to Aβ42 monomers and fibrils. (**A**) Principle of diffusion-based sizing of Alexa647-labelled species (purple) in the presence of a larger interaction partner (orange). The line in the middle of the channel does not exist in the device, but indicates how the laminar flow becomes separated into non-diffused and diffused half, respectively, for detection at the end of the channel. (**B**) Size of a full-length IgG (1igy.pdb) compared with Aβ42 fibril (5kk3.pdb): 9 planes of one filament are viewed from the side (top) and fibril end (bottom). Each plane contains two monomers in darker and paler colour, with buried epitope residues indicated in one monomer. Each double-headed arrow corresponds to 2.5 nm. Structural models were prepared using MOLMOL (26). (**C-F**) Diffused fraction of Alexa647-antibodies in the absence and presence of increasing concentrations of unlabeled Aβ1-42 fibrils, concentration given in monomer units. (**G-J**) Diffused fraction of Alexa647-Aβ(MC1-42) monomers in the absence and presence of increasing concentrations of each antibody. Examples of data for ^ch^aducanumab are in panels **C,G**; m266 **D,H**; 3D6 **E,I**; ^ch^gantenerumab **F,J**. Data for isotype control are in Fig. S12. All data were analysed by fitting directly to the observed diffused fraction of fluorescent species at the end of the microfluidic channel. The data in panels G-J were fitted assuming a fixed stoichiometry of *n=2* monomer binding sites per antibody, and with three adjustable parameters: *K*_*D*_ and the fractions of each state (free and bound) in the diffused fraction. The data in panels C-F were fitted with four adjustable parameters: *n* (=stoichiometry), *K*_*D*_ and the fraction of each state (free and bound) in the diffused fraction. (**K**) Antibody saturation versus free fibril site concentration, calculated from the fits in panels C-F and Fig. S12. (**L**) Monomer saturation versus free antibody concentration, calculated from the fits in panels G-J and Fig. S12. (**M**) Fraction of fibrils with at least one antibody bound calculated using the fitted values of K_D_ and *n*. (**N**) Summary of obtained affinities of each antibody for fibrils (pKDfibril) *versus* monomers (pKD monomer).

The microfluidic diffusional sizing measurements further reveal a moderate affinity of ^ch^gantenerumab for Aβ42 monomers (pK_D_ = 6.3±0.1, KD=485±100 nM) and a higher affinity for fibrils (pK_D_ = 7.5±0.3, KD=30±15 nM). The stoichiometry of one ^ch^gantenerumab per 43±4 monomers in the fibril, was obtained using short fibrils (50 nm, i.e. ca. 100 planes per filament), thus suggesting that ^ch^gantenerumab binds favorably to fibrils ends, rationalizing the specific kinetic fingerprint of this antibody in inhibiting fibril growth (Fig. 1). These findings are consistent with Aβ residues 18-27 forming an essential part of the epitope (9,25); several side-chains of this segment are buried in the fibril core and only accessible at the fibril ends (22,23; Fig. 3B). The affinities obtained here in solution are of a similar order of magnitude but generally lower than those reported for interactions with gantenerumab at surfaces (24), an observation which can be typical of surface-based methods that can overestimate binding affinities, which could also be influenced by the labelling or differences between the human and mouse versions of this antibody. Next, for 3D6 we find a relatively high affinity for Aβ42 monomers (pK_D_ = 7.4±0.1, KD=38±8 nM) and a very high affinity for fibrils (pK_D_ = 9.57±0.36, KD = 0.27±0.20 nM) with a stoichiometry of 1 antibody per 47±10 monomers in the fibril. All kinetic data with 3D6 can be explained by a reduction of k_+_ (Fig. 1C, S6C, S8E). Finally, for m266 we observe no measurable binding to fibrils but a very high affinity for monomers (pK_D_ = 8.5±0.3, KD = 3±2 nM), which is consistent with suppression of primary nucleation (Fig. 1) and the very large concentration of Aβ42 eluting in early fractions likely representing antibody-bound monomers (24, Fig. S10).

Taken together, the kinetic and binding data suggest that the antibodies studied here have dramatically different modes of action. In particular, the kinetic fingerprints shown in Fig 4a reveal that only ^ch^aducanumab specifically reduces the rate of the secondary nucleation pathway. To verify this mode of action, we benchmarked the activity of ^ch^aducanumab against the effect of a known and highly specific secondary nucleation inhibitor, the chaperone domain Brichos from pro-SPC (10). The data shown in Fig 4a reveal that ^ch^aducanumab has the same kinetic fingerprint as Brichos while the other antibodies exhibit a different type of behavior. To demonstrate the highly specific nature of the inhibition of secondary nucleation by ^ch^aducanumab, we next investigated the combined effect of the antibodies together with the chaperone. Since the kinetic fingerprints show that both ^ch^aducanumab and the chaperone inhibit secondary nucleation specifically, the analysis predicts that no effects on the aggregation kinetics will be observed upon the addition of ^ch^aducanumab in the presence of excess Brichos. By contrast, since the kinetic fingerprints for gantenerumab and 3D6 show that these antibodies inhibit processes other than secondary nucleation, an additive inhibitory effect is expected even in the presence of excess chaperone. These predictions are verified by the data probing the aggregation process in the presence of each of the antibodies and Brichos (Fig 4b), demonstrating the specific activity of ^ch^aducanumab in inhibiting the secondary nucleation pathway.

**Figure 4.**
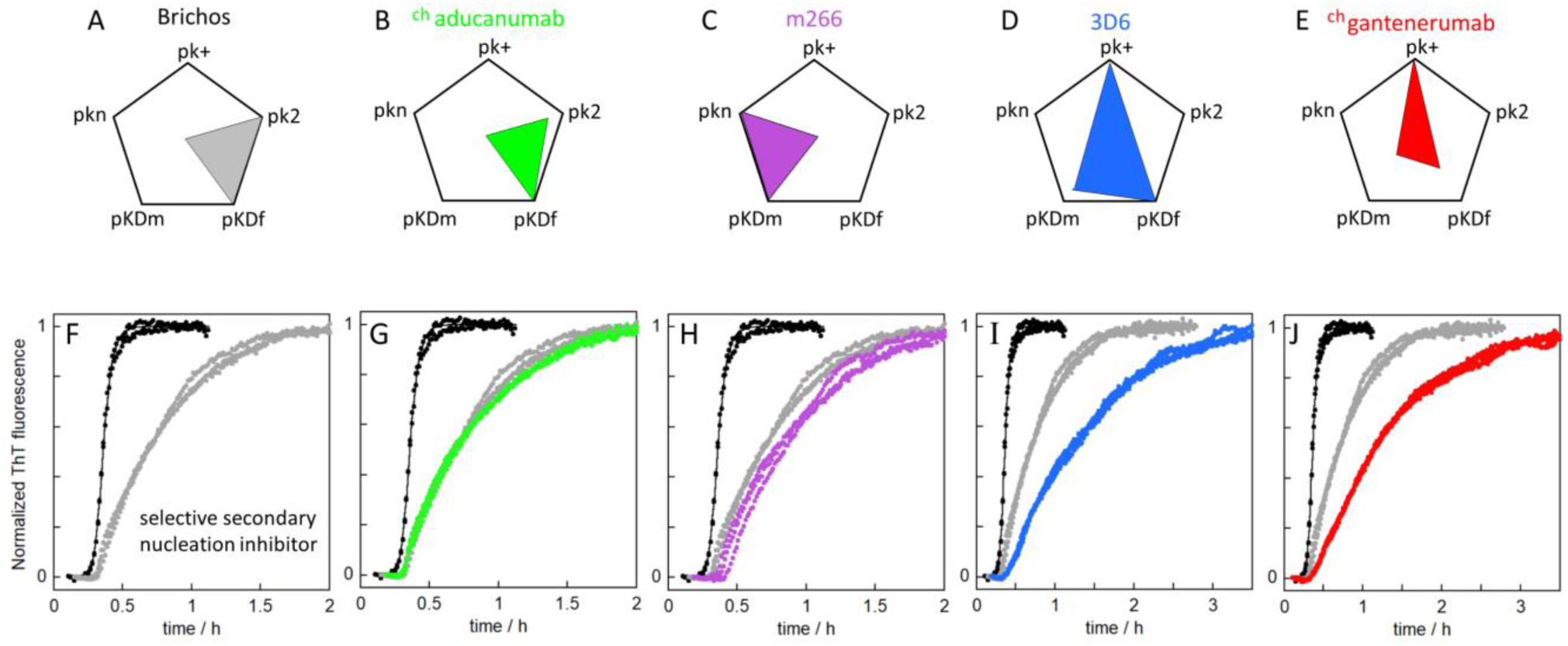
Kinetic fingerprints differentiate the mechanism of action of the anti-Aβ antibodies. **(A-E)** Effects of pro-SPC-Brichos (**A**), ^ch^aducanumab (**B**), m266 (**C**), 3D6 (**D**) and ^ch^gantenerumab (**E**) on the microscopic rate constants pictured in a pentagon together with the pK_D_ values for monomers (pK_Dm_) and fibrils (pK_Df_). The centre of each pentagon corresponds to unperturbed rate constants and a high pk value signifies a large reduction in rate constant. (**F-J**) Aggregation kinetics of 3 µM Aβ42 in 20 mM HEPES, 1 mM CaCl_2_, 150 mM NaCl, pH 8.0 in the absence (black) and presence of 3 µM pro-SPC-Brichos (grey, **F**), 3 µM pro-SPC-Brichos and 250 nM ^ch^aducanumab (green, **G**), 3 µM pro-SPC-Brichos and 250 nM m266 (purple, **H**), 3 µM pro-SPC-Brichos and 250 nM 3D6 (blue, **I**), 3 µM pro-SPC-Brichos and 250 nM ^ch^gantenerumab (red, **J**); in (G-J) the aggregation kinetics are compared with those in the presence of Brichos (grey).

In addition to direct inhibition of secondary nucleation, a second potential mechanism of action to reduce oligomer flux is the removal of the reactant species required for this process, monomers and fibrils. Monomer removal is likely to be challenging *in vivo* since it would require stoichiometric amounts of antibody to enter the brain, but crucially removal of the catalytic fibril surface is possible at sub-stoichiometric antibody concentrations. This latter process requires a significant fibril-bound antibody population; the binding analysis reveals that ^ch^aducanumab has a KD for fibril around 1 nM, and is significantly more specific to aggregated species than the other clinical antibodies tested here (Fig. 3N). Indeed, calculating the fraction of fibrils that are antibody bound as a function of free antibody concentration predicts that ^ch^aducanumab will be highly fibril bound even at low concentrations, whereas 3D6 in spite of its higher affinity, and especially ^ch^gantenerumab, and due to its lower stoichiometry per fibril, require relatively higher concentrations of free antibody to achieve the same fraction of bound fibrils (Fig. 3M).

Our results demonstrate that kinetic fingerprints acquired *in vitro* (Fig. 4) are able to reveal the strikingly varied mechanisms of action of clinical stage antibodies that have commonly been classified together as a single category of Aβ-targeting therapies. We illustrate the power of this approach by demonstrating that ^ch^aducanumab is effective at both reducing the rates of secondary nucleation, leading to a decrease in the flux of oligomeric forms of Aβ, as well as binding amyloid fibrils, thus targeting them for removal, a unique mechanism of action that may help explain its reported clinical activity (8, 27).

**Supplementary Information** is available for this paper.

## AUTHOR CONTRIBUTIONS

SL, OH designed the study. SL expressed and purified recombinant Aβ1-42 and Aβ(MC1-42) peptides. SL, ML and ES developed and optimized the protocol for recombinant Aβ1-42 production and purification. FQ, TO’M, TB and PHW generated and purified the antibodies. OH provided the pooled CSF samples. SL isolated Aβ42 monomers and antibodies by SEC and performed the aggregation kinetics experiments before unblinding. SL performed kinetic analysis of the data before unblinding. GM, SC, TPJK performed additional kinetic analysis of the data after unblinding. SL produced and purified the Alexa647-labelled forms of Aβ42 and all antibodies. SL performed the oligomer quantification experiments, and KB performed all mass spectrometry analyses before unblinding. TS performed the diffusion measurements of K_D_ and stoichiometry before unblinding. SL, TS, SD, CKX and GM analyzed the diffusion data. SL, MV, SC, TPJK, OH wrote the paper with input from all co-authors.

## DATA AVAILABILITY

The data that support the findings of this study are available from the corresponding author upon reasonable request.

## Supplementary information

**Figure S1.**
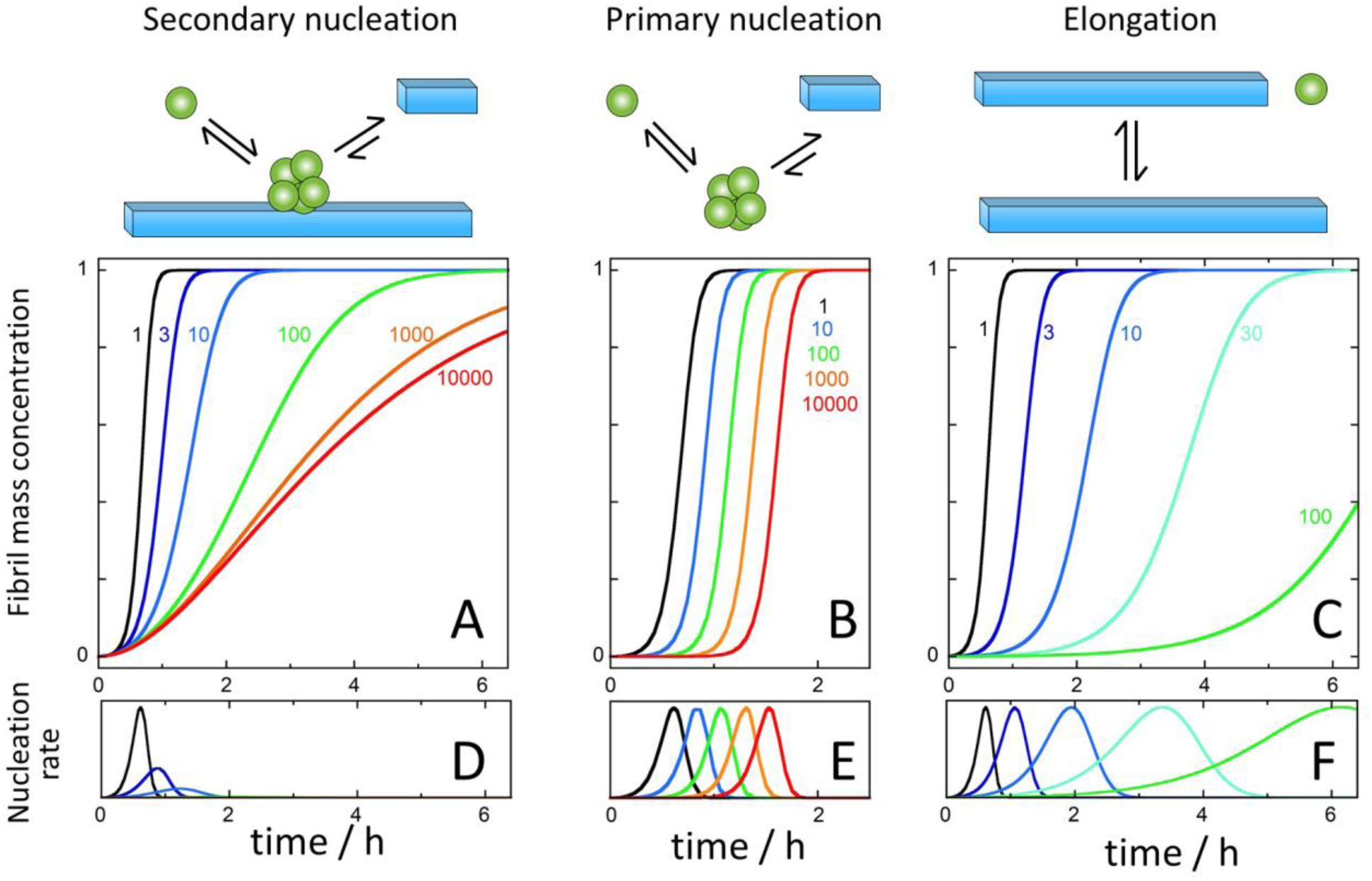
Theoretical aggregation curves and nucleation rates. (**A-C)** Predicted changes to the macroscopic aggregation curve if the rate constant for one specific microscopic step is selectively reduced by a factor of 3 (marine blue), 10 (blue), 30 (cyan), 100 (green), 1000 (orange) or 10000 (red). The reference curve is calculated for 3 µM Aβ42 in 20 mM HEPES, 140 mM NaCl, 1 mM CaCl_2_ pH 8.0 (*18*) with k_n_k_+_ = 2.5.10^−3^ M^-2^s^-2^, k_2_k_+_ = 2.1.10^13^ M^-3^s^-2^, √K_M_ = 0.25 µM; n_c_ = 1, n_2_ = 2. (**A-C)** Selective inhibition of: secondary nucleation **(A)**, primary nucleation **(B)**, and elongation **(C)**. The total nucleation rate underlying each trace in panel **A, B** and **C** is shown in panel **D, E**, and **F**, respectively. Note that since the macroscopic curves are sensitive to the rate constant products (k_n_k_+_ and k_2_k_+_) precisely the same effect as displayed in panel **C** can in principle arise if both k_n_ and k_2_ are reduced in parallel by the same factor.

**Figure S2.**
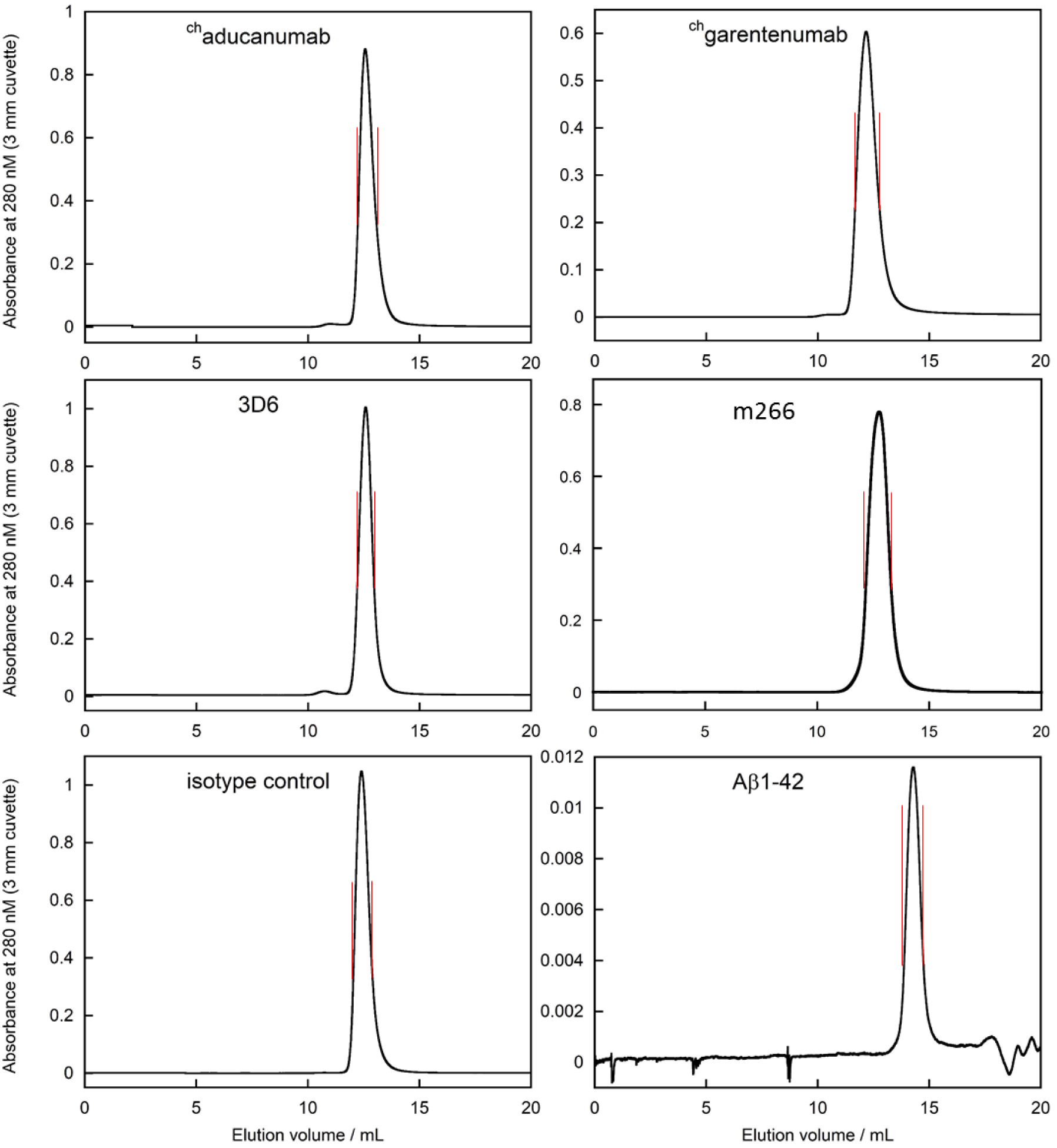
Isolation of monomeric Aβ42 and antibodies by size exclusion chromatography. Isolation of the antibodies on a 1 × 30 cm Superdex 200 column (GE Healthcare) in 20 mM HEPES/NaOH, 140 mM NaCl, 1 mM CaCl_2_, pH 8.0 just prior to the kinetics experiment (first five panels). Isolation of recombinant Aβ1-42 on a 1 × 30 cm Superdex 75 column (GE Healthcare) in 20 mM HEPES/NaOH, 1 mM CaCl_2_, pH 8.0 just prior to the kinetics experiment (last panel). 140 mM NaCl was added from a 4.2 M NaCl stock after size exclusion to limit losses on the column. In each case, the fraction between the vertical red lines was collected.

**Figure S3.**
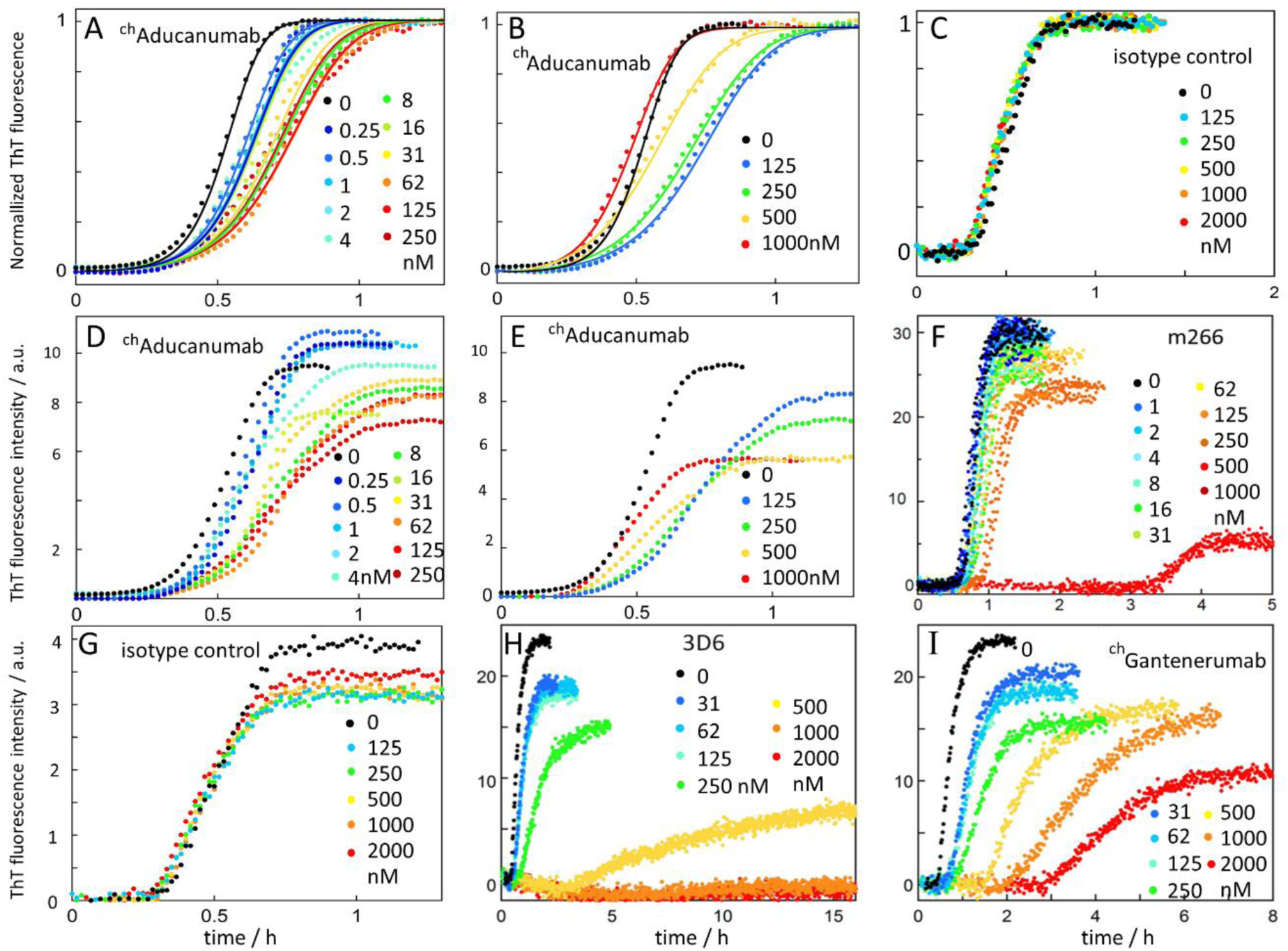
Aggregation kinetics data of Aβ1-42 in the absence and presence of antibodies. All panels show data acquired using recombinant Aβ1-42 at 37 °C under quiescent conditions in 20 mM HEPES/NaOH, 140 mM NaCl, 1 mM CaCl_2_, pH 8.0 in PEGylated plates. (**A-C**) Normalized data with ^ch^aducanumab (**A,B**) and isotype control (**C**). The same data are shown in non-normalized form in panels (**D,E**) and (**G**), respectively. Panels (**F, H, I**) show the same data as shown in Fig. 1, and in addition panel **H** includes data at 500-2000 nM 3D6. The fitted lines in panel A are with reduced values of k_2_. The fits in panel B, to normalized data in the absence and presence 250 -1000 nM ^ch^aducanumab, allow for a decrease in k_2_ and an increase in k_n_.

We note that at very high concentrations (>500 nM), which are unlikely to be realized *in vivo*, ^ch^aducanumab accelerates the bulk Aβ42 aggregation process by increasing the primary nucleation rate. Indeed, the data at 250-1000 nM ^ch^aducanumab are best fitted assuming that the effect on k_2_ is combined with an increase in k_n_ (Fig. S4). Interestingly, at high concentrations of each antibody, the ThT intensity is significantly reduced, which for m266 and 3D6, is found to be related to a significant concentration of Aβ1-42 remaining in solution (Fig. S5), implying the possibility of monomer binding which we assess directly in solution (Fig. 3).

**Figure S4.**
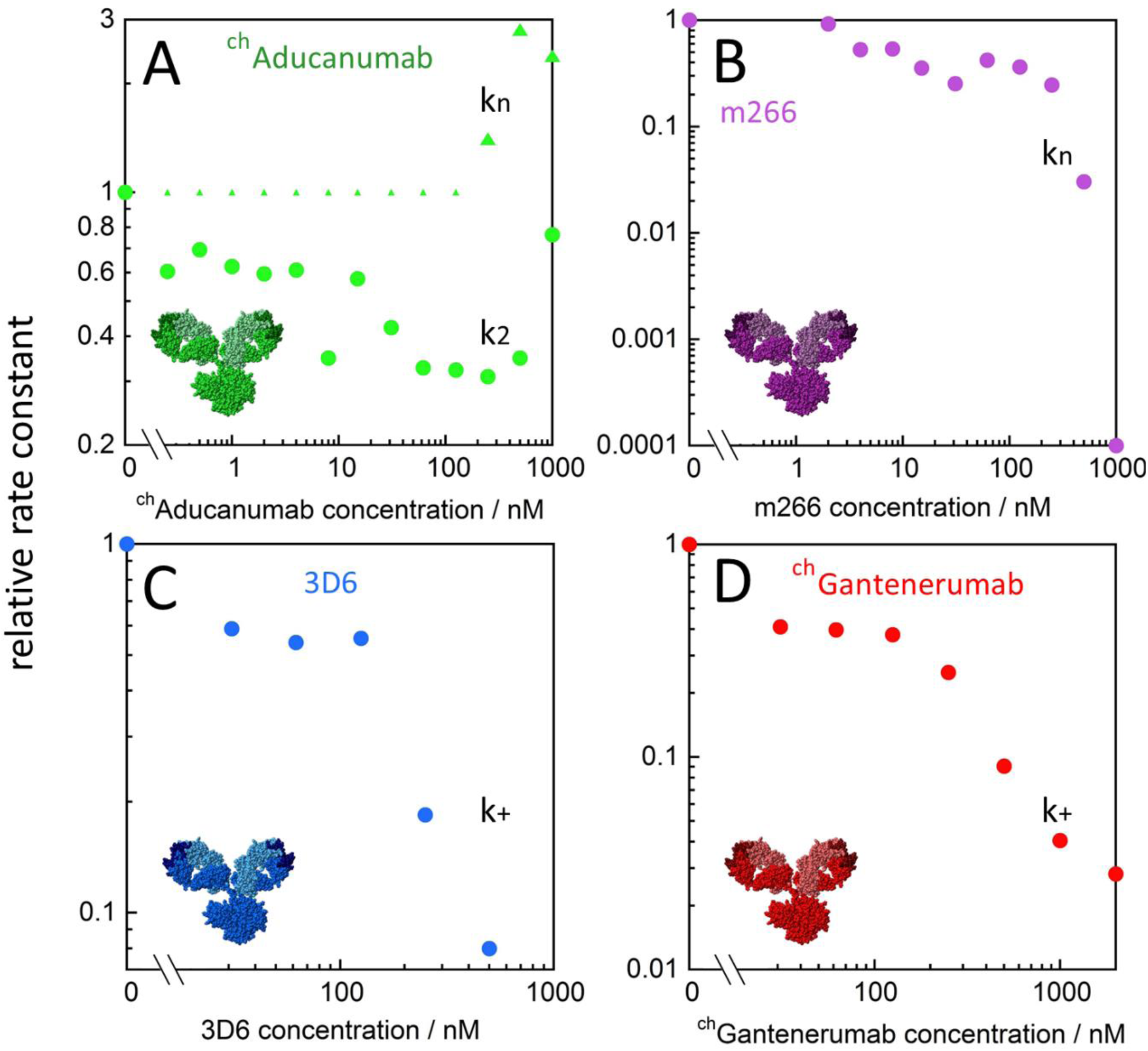
Effects of the antibodies on the rate constants. The rate constant values relative to Aβ42 alone are shown as a function of the antibody concentration for the case that produced the best fit to the data for each antibody (see Fig. 1). For ^ch^aducanumab (**A**) we thus show the effect on the rate constant for secondary nucleation, k_2_, up to 125 nM antibody, and thereafter the effects in k_n_ and k_2_. For m266 (**B**) we show the effect on the rate constant for primary nucleation, k_n_. For 3D6 (**C**) and ^ch^gantenerumab (**D**) we show the effect on the elongation rate constant, k_+_.

**Figure S5.**
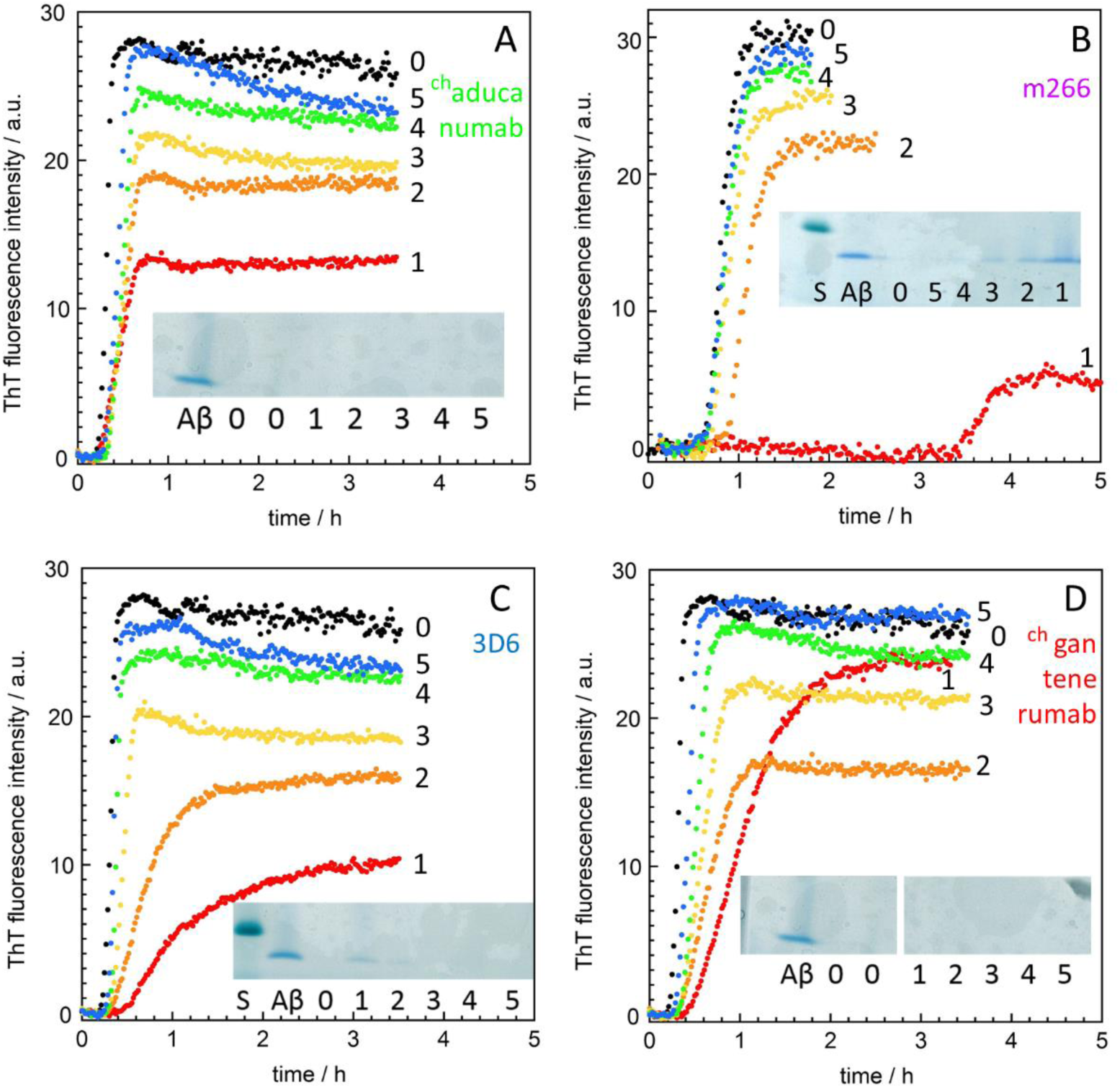
Measurements of the amounts of Aβ42 monomers remaining in solution at the end of the aggregation reaction. We measured the Aβ1-42 monomers remaining in solution after reaching the ThT plateau in the presence of antibodies that cause a drop in final ThT intensity. Sample from multiple wells (300-400 µl) were collected after reaching the ThT plateau and fibrils were removed by centrifugation. A small sample (10 µl) from the top of the supernatant was analyzed using SDS PAGE (the standard, S, in panel C and B is at 10 kDa). Lane Aβ shows monomers before aggregation, the lane labeled 0 shows the remaining monomer at the plateau for Aβ1-42 alone. Lanes 1-5 correspond to 1:1 serial dilutions of antibody from 500 nM (^ch^aducanumab, 3D6 and ^ch^gantenerumab) or 1 µM (m266). Remaining Aβ1-42 monomers at the plateau are clearly detected with 1 µM, 500 nM and 250 nM m266, and with 500 nM and 250 nM 3D6. Faint Aβ1-42 bands are seen with 125 nM m266 and 125 nM 3D6. Thus, the reduced ThT intensity at the plateau intensity is, at least in part, due to less monomer being consumed in the reaction when high concentration of m266 or 3D6 are present. No Aβ1-42 monomer is detected by this method at the plateau in the presence of ^ch^aducanumab or ^ch^gantenerumab, indicating that the effect on the ThT plateau intensity is not due to less monomer being consumed in the reaction. Instead, it is likely an effect of these antibodies interfering with the ThT signal. All data were acquired using Aβ1-42 at 37 °C under quiescent conditions in 20 mM HEPES/NaOH, 140 mM NaCl, 1 mM CaCl_2_, pH 8.0 in PEGylated plates.

**Figure S6.**
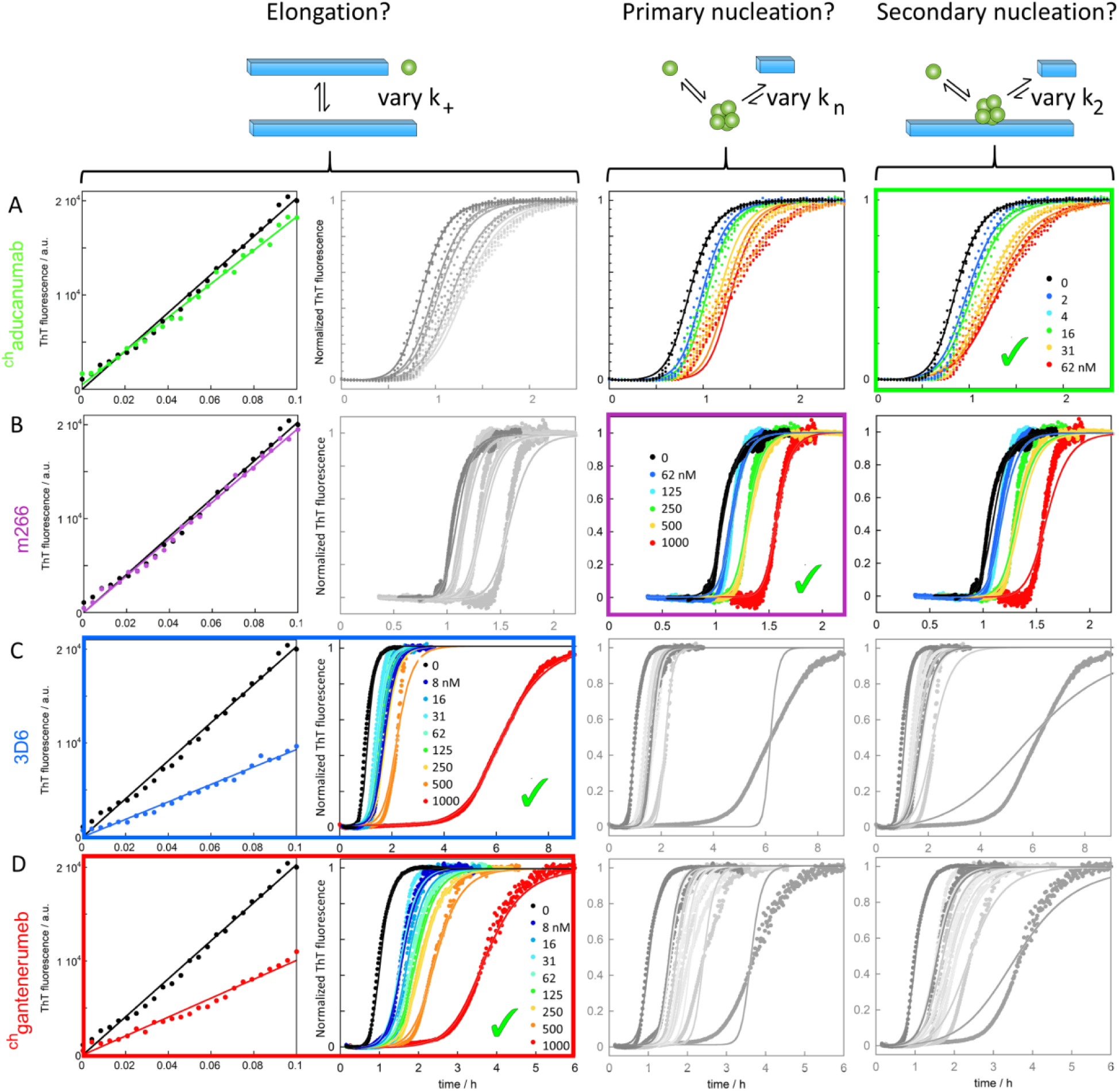
Effects of the antibodies on the aggregation kinetics of Aβ1-42 in CSF. ThT fluorescence as a function of time for reactions starting from 6 µM Aβ1-42 in 66% CSF, 20 mM HEPES/NaOH, 140 mM NaCl, 1 mM CaCl_2_, pH 8.0 in the absence and presence of ^ch^aducanumab (A), m266 (B), 3D6 (C) or ^ch^gantenerumab (D). The colour codes for the antibody concentrations are given in nM in each panel. The left column shows data obtained in the presence of 30% preformed seeds, with linear fits, and the following three columns show non-seeded data fitted three times with the selective variation of one rate constant in each column. For ^ch^aducanumab, the heavy seeded data rule out an effect on k_+_, and we can identify a reduction in k_2_ as the model which best fits the ^ch^aducanumab data in CSF; this is verified using light seeding in CSF (Fig. S7). Likewise, for m266 the heavy seeded data rule out an effect on k_+_, and we can identify a reduction in k_n_ as the model which best fits the m266 data in CSF. For 3D6 as well as ^ch^gantenerumab, the seeded data show an effect on k_+_, which indeed explains also the non-seeded data in CSF. Note that the x-axis covers 2.2, 6 or 8 h depending on the magnitude of the effect of each antibody.

**Figure S7.**
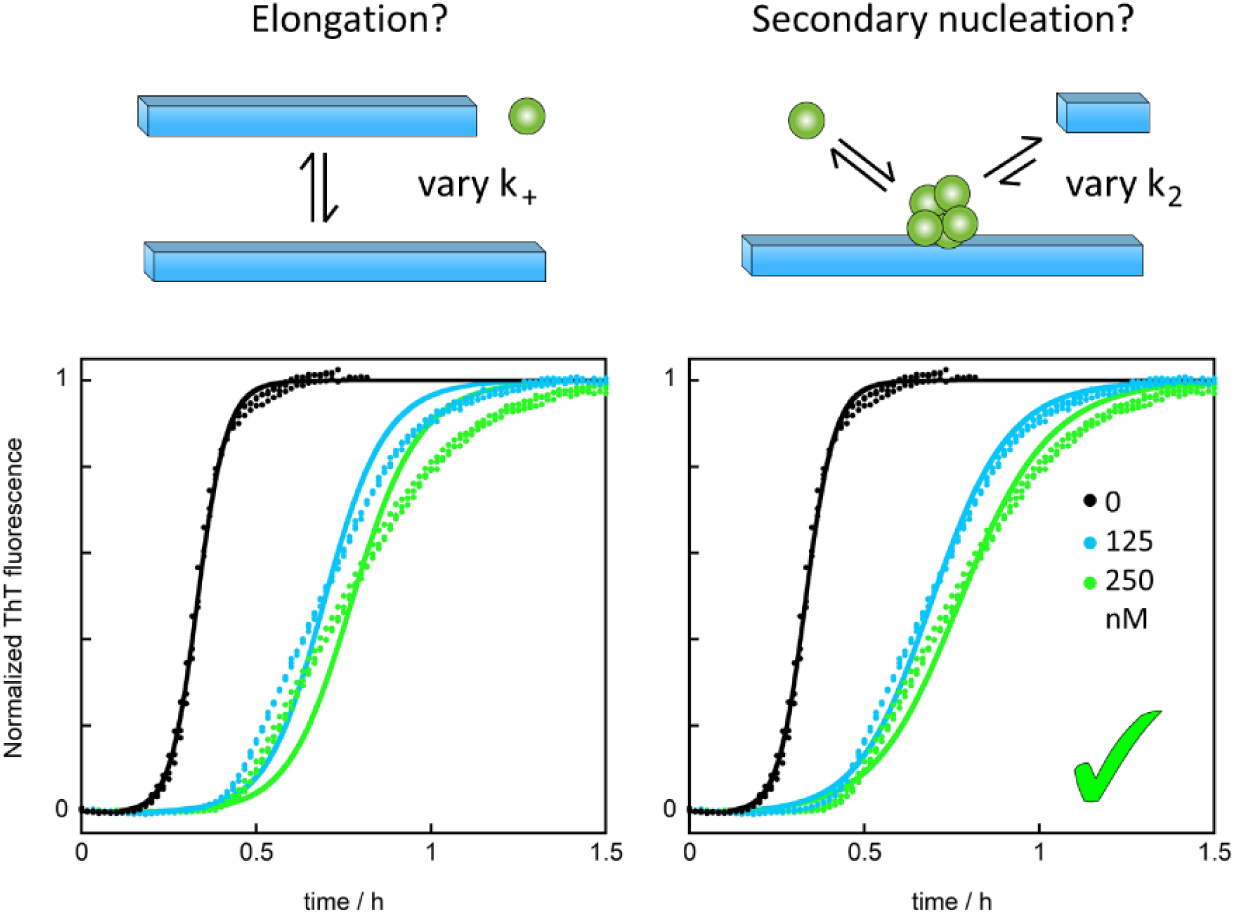
Effects of chaducanumab on light seeded (2%) aggregation kinetics of Aβ1-42 in CSF. ThT fluorescence as a function of time for reactions starting from 6 µM Aβ1-42 with 120 nM preformed seed fibrils in 66% CSF, 20 mM HEPES/NaOH, 140 mM NaCl, 1 mM CaCl_2_, pH 8.0 in the absence and presence of ^ch^aducanumab at 125 and 250 nM. The fits in the left panel assumes a constant value of k_2_ and curve-specific values of k_+_. The fits in the right panel assumes a constant value of k_+_ and curve-specific values of k_2_. The inclusion of a low amount of seeds passes primary nucleation and makes the data independent of this step; these data thus validate a reduction of secondary nucleation rate as the primary role of ^ch^aducanumab in CSF.

**Figure S8.**
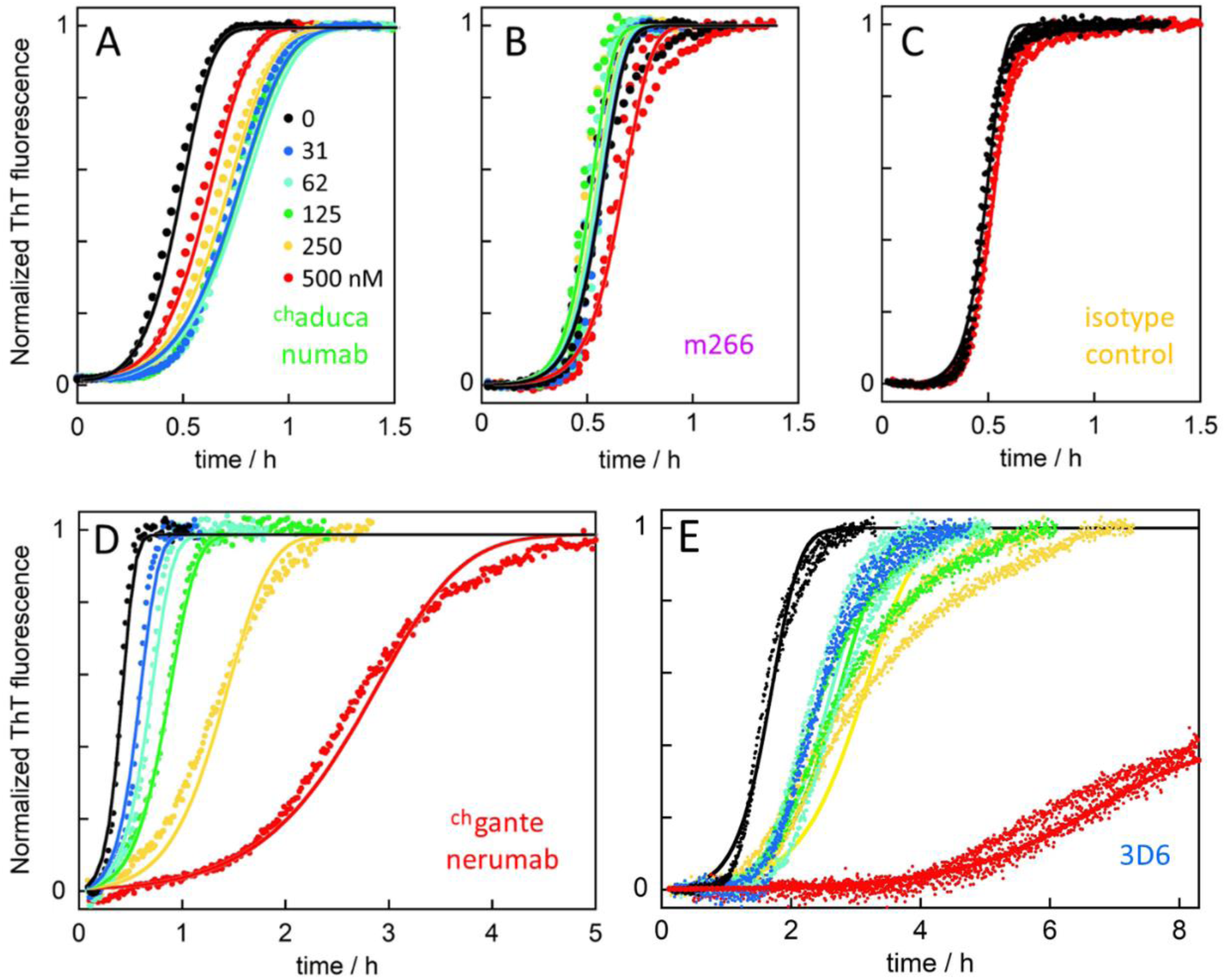
Aggregation kinetics in the presence of preformed seed fibrils at low concentration (2%). Normalized ThT fluorescence as a function of time for reactions starting from 3 to 4 µM Aβ1-42 monomer and 2% (60 to 80 nM) Aβ1-42 fibrils in 20 mM HEPES/NaOH, 140 mM NaCl, 1 mM CaCl_2_, pH 8.0 in the absence (black) and presence of A) ^ch^gantenerumab, B) m266, C) isotype control antibody (yellow), D) ^ch^aducanumab, or E) 3D6 at five concentrations as given by the colour code in panel A. The fitted curves in panels A-C allow only a variation of k_2_, whereas the fitted curves in panels D and E allow only a variation of k_+_.

**Figure S9.**
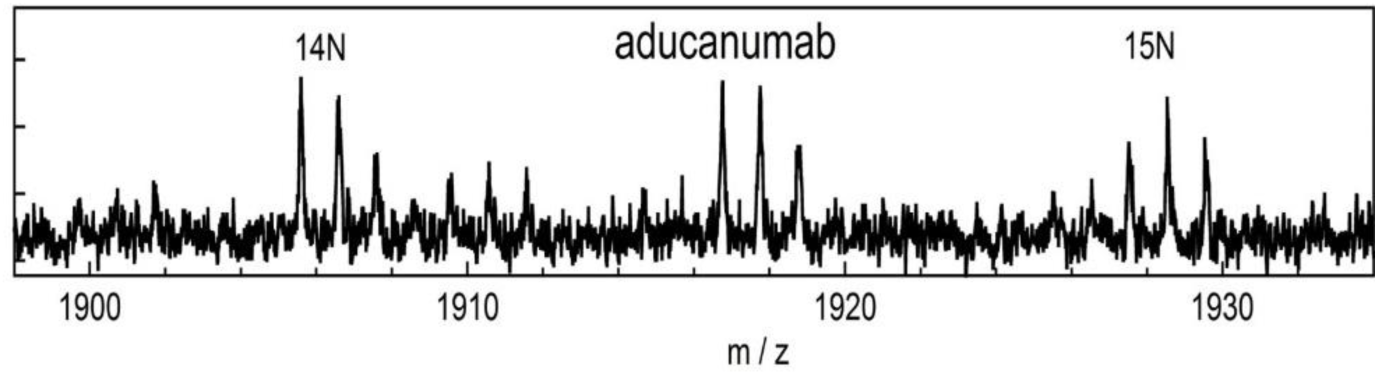
MALDI TOF/TOF analysis of Aβ1-42 in the presence of chaducanumab. Example of a MALDI TOF/TOF spectrum with direct spotting of samples collected at the half-time of aggregation of Aβ1-42 in the presence of ^ch^aducanumab.

**Figure S10.**
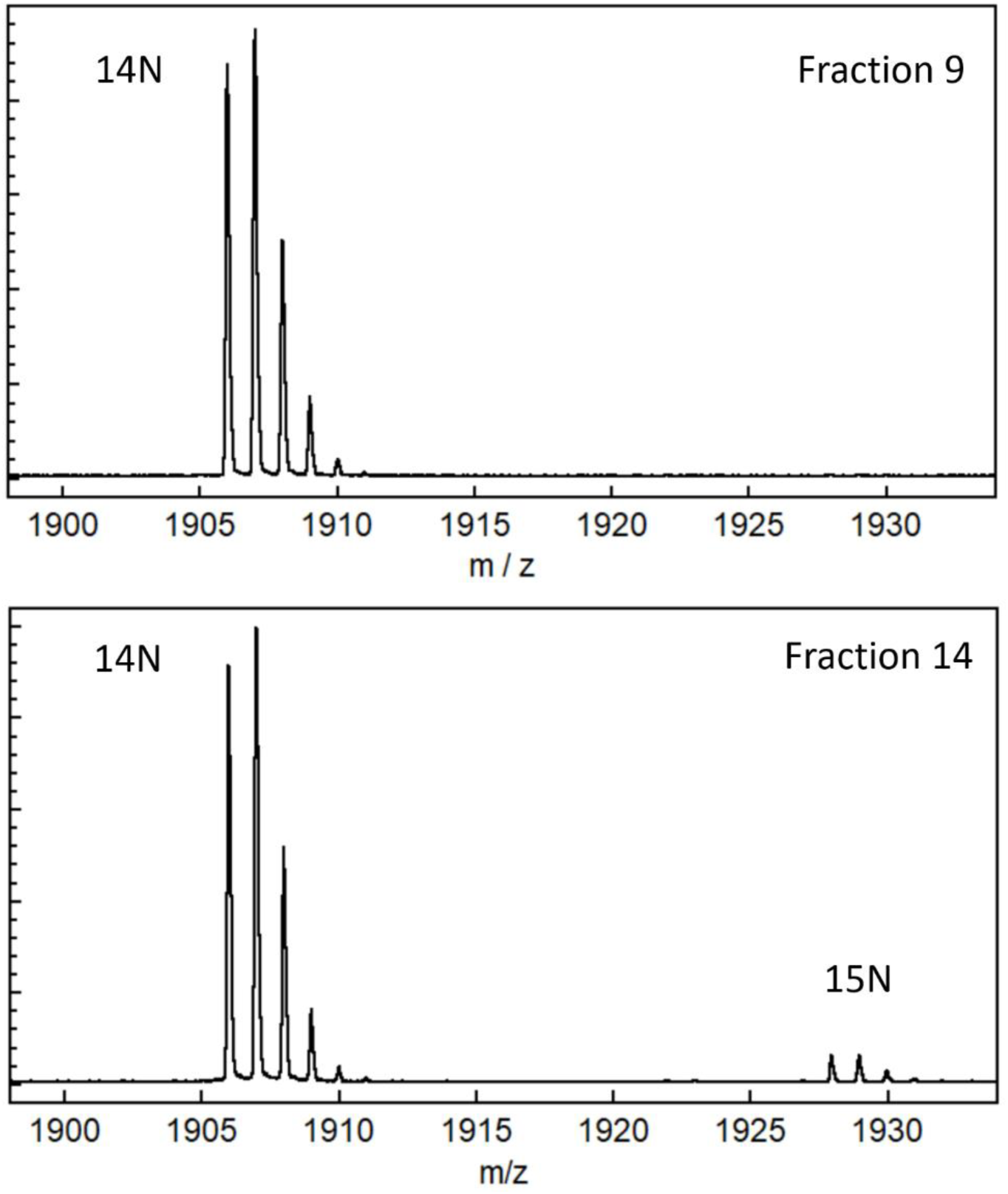
MALDI TOF/TOF analysis of Aβ1-42 in the presence of m266. Example of a MALDI TOF-TOF spectra with HPLC separation before spotting for samples collected at the half-time of aggregation of Aβ1-42 in the presence of m266. In Fraction 9 there is so much 14N peptide that the signal from 15N is suppressed.

**Figure S11.**
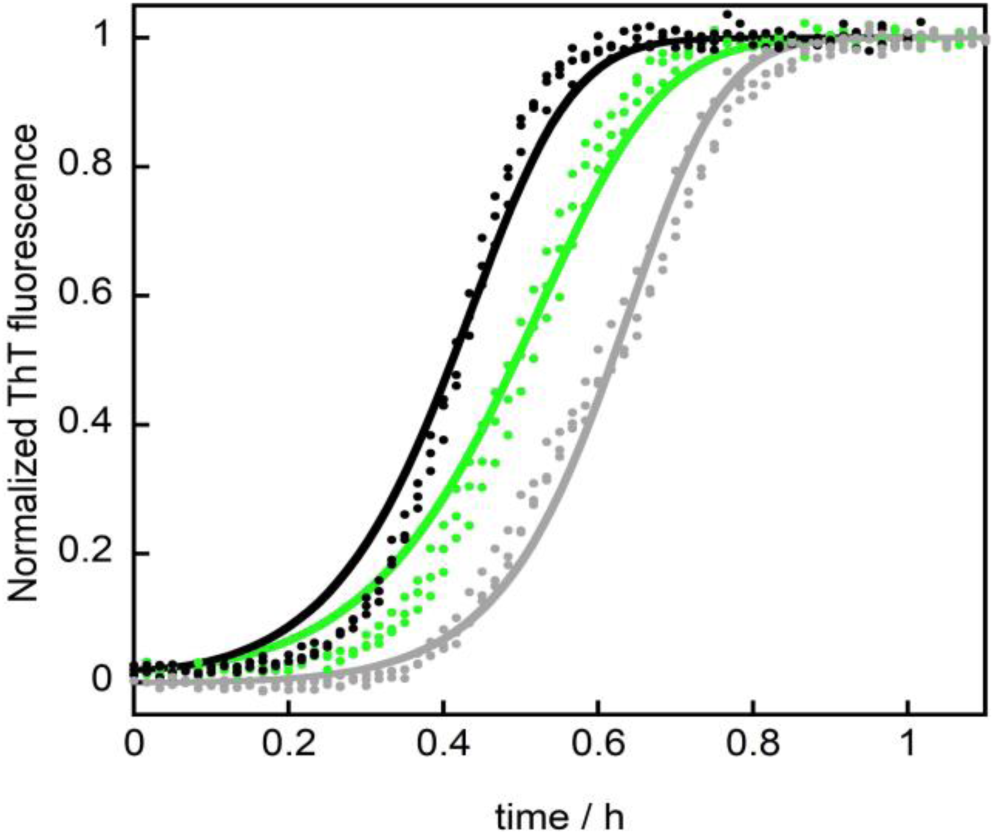
Effects of chaducanumab in light seeded (2%) aggregation kinetics of Aβ1-42. ThT fluorescence as a function of time for reactions starting from 3 μM Aβ1-42 in the absence (grey) or presence of 60 nM preformed seed fibrils in 20 mM HEPES/NaOH, 140 mM NaCl, 1 mM CaCl_2_, pH 8.0. The seeds were made in the absence (black) or presence of ^ch^aducanumab (green). The total concentration of ^ch^aducanumab in the samples for the data shown in green was 6 nM. The fits assume a constant value of k_+_ and curve-specific values of k_2_. The relative values of k_2_ obtained are 1.0 (grey), 1.0 (black) and 0.67 (green). Thus, forming the seeds in the presence of ^ch^aducanumab leads to a 33% reduction in apparent k_2_, very similar to what we observed with 6 nM ^ch^aducanumab in the non-seeded data (Fig S4). We conclude that a significant fraction of the inhibitory effect on secondary nucleation originates from interaction of the antibody with fibrillar aggregates.

**Figure S12.**
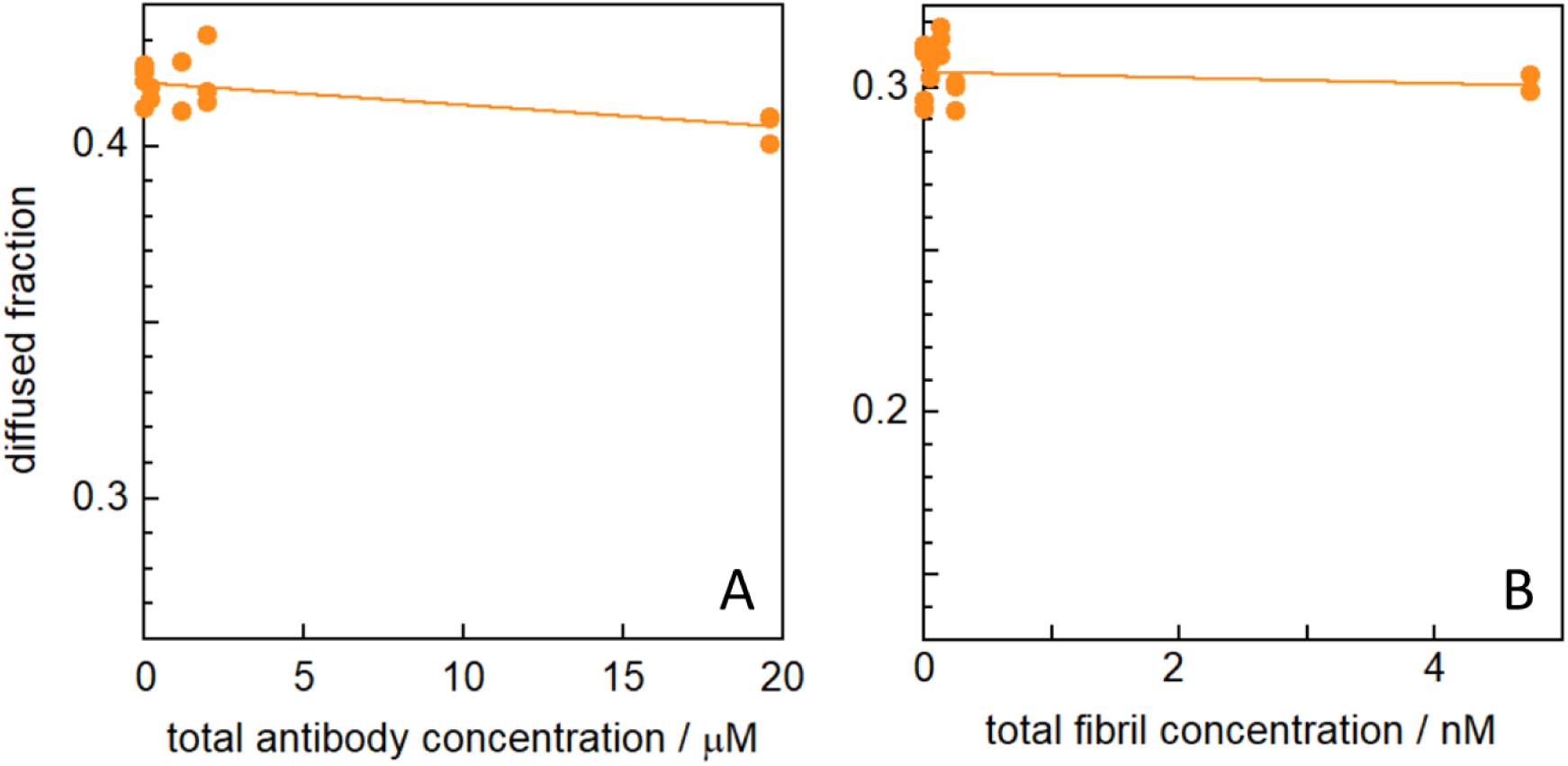
Antibody binding analysis in solution with isotype control antibody. **A**) Diffusion of Alexa-647-Aβ(MC1-42) in the absence and presence of increasing concentration of isotype control antibody. **B)** Diffusion of Alexa-647-labelled isotype control antibody in the absence and presence of increasing concentrations of unlabelled Aβ1-42 fibrils. In both panels are shown the fraction of fluorescence appearing in the diffused half at the channel outlet as a function of total concentration of the non-labelled species. The solid lines are fitted straight lines.

**Figure S13.**
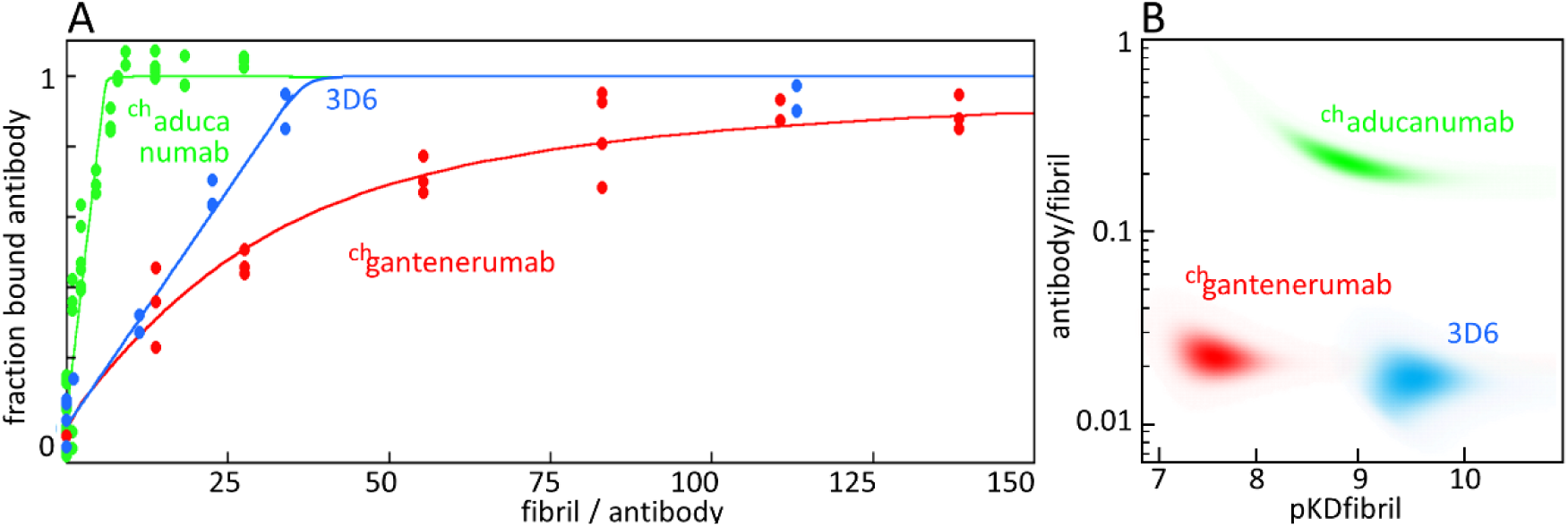
Antibody binding analysis in solution. **A)** The same data as in figure 3C,E,F replotted with the ratio of fibril to antibody concentration on the X-axis. The fibril concentration is in monomer units. **B)** Examples of results of Bayesian analysis.

## Online Methods

### 1.1 Chemicals and solutions

All chemicals were of analytical grade. All buffers were filtered (200 nm) and extensively degassed before use. Thioflavin T (ThT) was from CalBiochem and was prepared as a stock solution in water, which was filtered through 200 nm filter and the concentration determined using absorbance. CSF was collected from de-identified healthy donors, pooled and frozen as multiple identical aliquots.

### 1.2 Expression of Aβ1-42 peptide

Due to the requirement of an ATG start codon yielding an N-terminal Met residue, Aβ1-42 cannot be expressed “as is” and requires a fusion tag. We employed the self-cleavable tag nPro in the form of its EDDIE mutant (*28*). EDDIE-Aβ1-42 with the amino acid sequence MELNHFELLYKTSKQKPVGVEEPVYDTAGRPLFGNPSEVHPQSTLKLPHDRGEDDIETTLRDLPRKGDCRSGNHLGPVSGIYIKPGPVYYQDYTGPVYHRAPLEFFDETQFEETTKRIG RVTGSDGKLYHIYVEVDGEILLKQAKRGTPRTLKWTRNTTNCPLWVTSC<u>DAEFRHDSG YEVHHQKLVFFAEDVGSNKGAIIGLMVGGVVIA</u>, with the Aβ1-42 sequence underlined, was expressed from a Pet3a plasmid (purchased from Genscript, Piscataway, New Jersey) in *E. coli* BL21 DE3 PlysS star in overnight express medium (2.5 mM Na_2_HPO_4_, 2.5 mM KH_2_PO_4_, 12 mM (NH_4_)_2_SO_4_, 1 mM MgSO_4_, 0.1 g/L glucose, 0.4 g/L lactose, 1 g/L glycerol, 10 g/L NaCl, 10 g/L tryptone, 5 g/L Bacto yeast extract, 50 mg/L ampicillin, 30 mg/L chloramphenicol).

### 1.3 Purification of Aβ1-42 peptide

Cell pellet from 4 L was sonicated 5 times in 80 mL 10 mM Tris/HCl, 1 mM EDTA, pH 8.5 containing a trace of DNase, with centrifugation at 18,000 g for 7 min between sonications. The inclusion body pellet (hard and grey, ca. 5-10 mL) after the 5^th^ sonication was dissolved in 150 mL 10 M urea, 10 mM Tris, 1 mM EDTA, 1 mM DTT, pH 8.5 by sonication, grinding, and stirring. The resulting solution (ca. 9.4-9.7 M urea) was diluted with 150 mL 10 mM Tris, 1 mM EDTA, 1 mM DTT pH 8.5 (thus yielding ca. 4.8 M urea) and loaded onto 2 × 20 mL DEAE-sepharose FF columns (GE Healthcare) in tandem. The columns were washed with 100 mL 4 M urea, 10 mM Tris, 1 mM EDTA, 1 mM DTT pH 8.5, and eluted with a 0-0.4 M NaCl gradient in the same buffer. Fractions containing EDDIE-Aβ1-42 were diluted 15 times with 1 M Tris, 1 mM EDTA, 5 mM DTT, pH 7.9 in glass bottles and left at 4°C for 48 h, total volume 1.5 L. During this time EDDIE slowly folds leading to auto-cleavage and release of Aβ1-42. The solution was then dialyzed (in 3.5 kDa MW cutoff dialysis bags that had been boiled 4 times in Millipore water before use) three times against 10 L of 5 mM Tris/HCl, 0.5 mM EDTA, pH 8.5. After this, the solution was poured into a 2 L beaker and supplemented with 50 g Q-sepharose big beads (GE Healthcare, equilibrated in 10 mM Tris/HCl, 1 mM EDTA, pH 8.5) and incubated for 0.5 h in the cold room with stirring using a propeller stirrer. The beads were collected on a Büchner funnel and washed with 200 mL 10 mM Tris/HCl, 1 mM EDTA, pH 8.5, and eluted with 8 × 100 mL 10 mM Tris/HCl, 1 mM EDTA, pH 8.5, 75 mM NaCl. The eluted fractions were examined using SDS PAGE with Coomassie staining. Fractions dominated by Aβ1-42 monomer were lyophilized, dissolved in 10 mL 6 M GuHCl and isolated from residual *E. coli* proteins, aggregates and small molecule contaminants using size exclusion chromatography on a Superdex 75 26/600 column at 4°C. The eluted fractions were examined using UV absorbance, agarose gel electrophoresis and SDS PAGE with Coomassie staining. Fractions corresponding to the center of the Aβ42 monomer peak were pooled, lyophilized, dissolved in 10 mL 6 M GuHCl, 20 mM sodium phosphate, pH 8.5 and subjected to a second round of size exclusion chromatography on a Superdex 75 26/600 column at 4°C. Fractions corresponding to the centre of the Aβ42 monomer peak were pooled, aliquoted and lyophilized.

### 1.4 Expression and purification of Aβ(M1-42) peptide

Aβ(M1-42) was expressed from a PetSac (derivative of Pet3a) plasmid in *E. coli* BL21 DE3 PlysS star in M9 minimal medium with ^15^NH_4_Cl as the only nitrogen source and purified essentially as described (*7, 29*): Cell pellet from 4 L culture was sonicated in 80 mL 10 mM Tris/HCl, 1 mM EDTA, pH 8.5 (buffer A) using a sonicator tip (half horn, 50% duty cycle, maximum output, 30-90 s). This step was performed in a glass beaker surrounded by an ice-water slurry. Inclusion bodies were isolated by centrifugation at 4°C and 18,000 × g for 10 min. Two more rounds of sonication and centrifugation were performed. The inclusion bodies were dissolved in 80 mL buffer A containing 8 M urea and diluted 4-fold in buffer A. 40 mL DEAE cellulose (Whatman DE23, equilibrated in buffer A with 2 M urea) was added, and the slurry was left on ice for 30 min (with periodic stirring using a glass rod). The solution was removed, and the resin washed with buffer A in a Büchner funnel on a vacuum flask, followed by two washes with 100 mL buffer A and elution with 8 × 30 ml buffer A containing 75 mM NaCl. The ion exchange purification was performed in batch mode to avoid concentrating Aβ peptide on the resin. The eluted fractions were examined using SDS PAGE with Coomassie staining. Fractions dominated by Aβ42 monomer were lyophilized, dissolved in 10 mL 6 M GuHCl and isolated from residual *E. coli* proteins, aggregates and small molecule contaminants using size exclusion chromatography on a Superdex 75 26/600 column at 4 °C. The eluted fractions were examined using UV absorbance, agarose gel electrophoresis and SDS PAGE with Coomassie staining. Fractions corresponding to the center of the Aβ(M1-42) monomer peak were pooled lyophilized, dissolved in 10 mL 6 M GuHCl, 20 mM sodium phosphate, pH 8.5 and subjected to a second round of size exclusion chromatography on a Superdex 75 26/600 column at 4°C. Fractions corresponding to the centre of the Aβ42 monomer peak were pooled, aliquoted and lyophilized.

### 1.5 Generation ad purification of anti-Aβ antibodies

Murine chimeric IgG2a/kappa versions of aducanumab (^ch^aducanumab), gantenerumab (^ch^gantenerumab), bapineuzumab (3D6) and solanezumab (m266) were generated as described (*24, 30*). The variable domain amino acid sequences of bapineuzumab (*31*) and gantenerumab (*9*) were selected based on publicly available sequence information and used to generate murine chimeric analogs. The resultant antibodies contain the variable heavy (V_H_) and variable light (V_L_) domains of the specific antibody and murine IgG2a/kappa constant heavy and constant light domains. An mIgG2a isotype control antibody P1.17 was generated from a murine myeloma cell line (ATCC TIB-10). All antibodies were expressed in CHO cells and purified by protein-A-affinity followed by ion-exchange chromatography. Purified antibodies were frozen as multiple identical aliquots. Prior to setting up the kinetics experiment, these aliquots were further purified using size exclusion chromatography on a 10×300 mm Superdex 200 column eluted in 20 mM HEPES/NaOH, 140 mM NaCl,1 mM CaCl_2_, pH 8.0, or in 20 mM sodium phosphate, 0.2 mM EDTA, pH 8.0. Examples of chromatograms are shown in Fig. S2.

### 1.6 Blinding of antibodies

The studies were run blinded such that several antibodies that were studied in parallel and their identities were not known to the person setting up the experiments and analyzing the data. In the experiments with Aβ1-42 at physiological salt buffer and in CSF there were four antibodies (isotype control, ^ch^aducanumab, 3D6, ^ch^gantenerumab). After un-blinding, the effect of m266 was investigated in physiological salt buffer and CSF.

### 1.7 Aggregation kinetics experiments

Aliquots of purified lyophilized Aβ1-42 were dissolved in 1 mL 6 M GuHCl, 20 mM sodium phosphate, pH 8.5, and monomer isolated by gel filtration on a Superdex 75 10/300 column in 20 mM HEPES/NaOH, 1 mM CaCl_2_, pH 8.0. The gel filtration step removes traces of pre-existent aggregates and exchanges Aβ peptide into the buffer used in the fibril formation experiments. An example chromatogram is shown in Fig. S2. The peptide concentration was determined from the absorbance of the integrated peak area using ε_280_ = 1400 L mol^-1^cm^-1^ as calibrated using quantitative amino acid analysis. The monomer generated in this way was diluted with buffer and supplemented with thioflavinT (ThT, Calbiochem) at a final concentration of 10 μM from a 2 mM stock (in water, filtered through a 0.2 µm filter). For experiments with Aβ1-42 at physiological ionic strength, 140 mM NaCl was added after gel filtration from a 4.2 M stock (in 20 mM HEPES/NaOH, 1 mM CaCl_2_, pH 8.0, filtered through a 0.2 µm filter). For experiments in the presence of CSF, 66% CSF was added from a 100% stock (buffered to pH 8.0 using saturated HEPES solution). The solution with maximum antibody concentration was mixed with Aβ42 solution so as to achieve the same Aβ42, buffer, salt, CaCl_2_ or EDTA, ThT and in some cases also CSF concentration as in the antibody-free sample. The antibody-free solution was then mixed in different proportions with the solution with maximum antibody concentration to achieve a dilution series with varying antibody concentration, keeping all other component concentrations constant. All samples were prepared in low-binding tubes (Axygen, California, USA) on ice, using careful pipetting to avoid introduction of air bubbles. Each sample was pipetted into wells of a 96 well half-area plate of PEGylated black polystyrene with a clear bottom (Corning 3881, Massachusetts, USA), 80 μL per well, 3 to 6 replicates of each solution in each plate. The final Aβ42 concentration was between 3 and 4 µM in the kinetic experiments and was constant within each plate. The kinetic assays were initiated by placing the 96-well plate at 37°C under quiescent conditions in a plate reader (FLUOstar Omega or FLUOstar Optima BMGLabtech, Offenburg, Germany). The ThT fluorescence was measured through the bottom of the plate every 60 s with a 440 nm excitation filter and a 480 nm emission filter. The whole setup with 3-6 replicates per concentration, was repeated at least three times for each combination of antibody, peptide, and buffer or CSF. Data shown in each panel are averages over all replicates in each run.

### 1.8 Seeded aggregation kinetics

The seed fibrils were prepared from 5 µM Aβ1-42 monomer solutions (in 20 mM HEPES/NaOH, 1 mM CaCl_2_, 140 mM NaCl, pH 8.0, 10 µM ThT, as above) by incubation at 37°C under quiescent conditions in a plate reader monitoring ThT fluorescence and collecting the fibrils when the fluorescence reached the post-transition plateau. Fresh Aβ1-42 monomer solutions were prepared as above. The seed fibrils were diluted to two times the desired final concentration and mixed with equal volume of fresh monomer solution to achieve a final seed concentration of either 30% or 2% of monomer concentration equivalent. Seed-free samples were made for comparison; the monomer solution was mixed with equivalent volume of buffer. Kinetic measurements were made as described above (1.7).

### 1.9 Aggregation kinetics in the presence of Brichos

The Brichos domains from pro-SPC was expressed and purified as previously described (*5, 31, 32*). Aggregation kinetics were followed as above under quiescent conditions in 20 mM HEPES/NaOH, 1 mM CaCl_2_, 140 mM NaCl, pH 8.0, 10 µM ThT, in PEGylated plates, for reactions containing 3 µM Aβ1-42 monomer, or 3 µM Aβ1-42 monomer plus 3 µM Brichos, or 3 µM Aβ1-42 monomer plus 3 µM Brichos plus 250 nM of one of the following antibodies: ^ch^aducanumab, m266, 3D6 or ^ch^gantenerumab.

### 1.10 SDS PAGE analysis of monomer concentration at the end of the reaction

This experiment was aimed at measuring if there is any remaining Aβ1-42 monomer at the ThT plateau in the absence and presence of the antibodies. Samples were collected at the end of the aggregation kinetics, fibrils were removed by centrifugation at 31,500 × g for 2 minutes in a benchtop centrifuge (Hettich Zentrifugen MIKRO 220R) and the supernatant was mixed 1:1 with SDS Loading buffer and analyzed using SDS PAGE on 4-20% polyacrylamide gradient gels (Novex) with a Tris/Bis-Tris Buffer system containing 0.1% SDS. The gels were stained with Coomassie Blue and the intensity of gel bands was compared to that of a monomer standard.

### 1.11 Kinetic analysis

The aggregation data in the absence and presence of different concentrations of antibodies were normalized and fitted using the Amylofit on-line platform (*16*) using the following integrated rate law for the normalized aggregate concentration

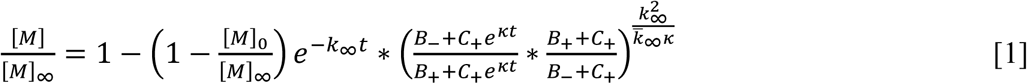

where the parameters are defined as follows

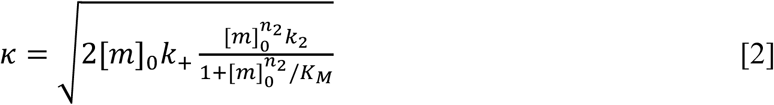

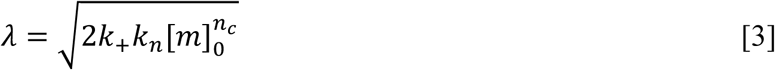

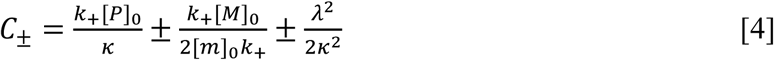

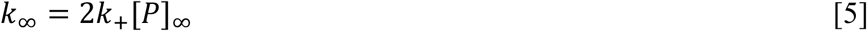

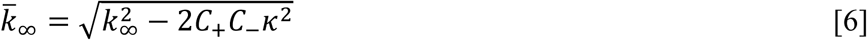

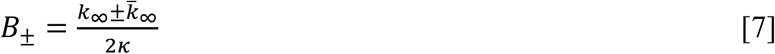

In these relations, [m]_0_ is the initial monomer concentration, [P]_0_ is aggregate number at the start of the reaction, [P]_∞_ is the aggregate number at equilibrium, that is, after reaction completion, [M]_0_ is the mass concentration of fibrils at the start of the reaction and [M]_∞_ is the mass concentration of fibrils at equilibrium. *k*_n_, *k*_2_, *k*_+_ are the rate constants for primary nucleation, secondary nucleation and elongation respectively. K_M_ is the saturation constant for secondary nucleation and *n*_*c*_ and *n*_*2*_ are the monomer scalings of primary and secondary nucleation, respectively. The values n_c_ = 0.0001, and n_2_ = 2 were used for fitting of all aggregation kinetics in 20 mM HEPES/NaOH, 1 mM CaCl_2_, 140 mM NaCl, pH 8 (*18*).

#### Microscopic mechanism of inhibition

The action of an inhibitor can be assessed through the analysis of macroscopic aggregation curves (*5, 6*). Inhibition of primary nucleation (rate constant k_n_) causes a lag-phase prolongation with no effect on the transition steepness, thereby delaying the generation of oligomeric species through secondary nucleation (Fig. S1B,E). This type of inhibition requires compounds that bind Aβ monomers, nuclei or other oligomers (*6,34-36*). By contrast, inhibition of secondary nucleation (with rate constant k_2_) causes little lag-phase extension but a reduction of the transition slope (Fig. S1A,D). This route for therapeutic intervention may significantly reduce toxicity (*5*), and requires inhibitors that block the catalytic surface of fibrils, or bind to on-pathway oligomers to prevent their conversion to fibrils (*5, 6, 34, 37, 38*). Finally, inhibition of elongation (with rate constant k_+_) causes both a lag-phase extension and a reduction in transition slope (*39*), an effect that may increase toxicity over time (*5*, Fig. S1C,F). When the ends of fibrils are blocked, a larger fraction of the monomers bind to their sides, where secondary nucleation and oligomer generation is catalyzed.

### 1.12 Oligomer size distribution and concentration

The concentration of oligomers and their size distribution was estimated at t_1/2_, the point in time where half the monomers were converted to fibrils (*14, 15, 19*). Aβ1-42 monomers were isolated by gel filtration in 20 mM HEPES/NaOH, 1 mM CaCl_2_, pH 8.0, followed by addition of 140 mM NaCl from a concentrated stock. Solutions containing 5 µM monomer and 100 nM antibody (isotype control, ^ch^aducanumab, ^ch^gantenerumab, m266 or 3D6) were prepared in the same buffer containing 1 µM thioflavin T. The reaction was followed in PEGylated plates (Corning 3881), 100 µl per well, by monitoring ThT fluorescence as described above. The reaction was stopped at t_1/2,_ i.e. when the ThT fluorescence reached half-way in between the starting baseline and the final plateau. Solution from 12 wells was combined and centrifuged for 2 minutes at 30,500 × g in a benchtop centrifuge (Hettich Zentrifugen MIKRO 220R). The top 1 mL of the supernatant was injected on a Superdex 75 10/300 column eluted in 20 mM ammonium acetate, pH 8.5, and 1 mL fractions were collected during elution. Each gel filtration fraction was lyophilized, dissolved in 10 µl water, supplemented with 1 pmol ^15^N-Aβ(M1-42) and subjected to digestion with AspN protease overnight. The sample was concentrated using a C18 zip tip and spotted (0.5 µl) on a steel plate, followed by application of 0.5 µl α-Cyano-4-hydroxycinnamic acid matrix solution and mass spectrometric analysis using an Autoflex Speed Matrix Assisted Laser Desortion Ionization (MALDI) Time-of-Flight (TOF)/TOF system (Bruker Daltonics). In cases of poor signal to noise or antibody fragment overlap, samples were separated by HPLC before spotting on the steel plate before analysis by MALDI-TOF/TOF or were analyzed by electrospray ionization (ESI) mass spectrometry using an Orbitrap instrument. The relative intensity of the peaks at 1906 (^14^N-Aβ7-22, DSGYEVHHQKLVFFAE) and 1928 (^15^N-Aβ7-22 isotope standard), was used to estimate the oligomer concentration in each fraction.

### 1.13 Alexa-647 labelling of peptides and antibodies

Aβ(MC1-42), a mutant with an extra Cys residue placed between the starting Met and Asp1 was used for fluorophore labelling (*40*). An aliquot of purified peptide monomer was dissolved in 6 M GuHCl, 10 mM DTT, pH 8.5, incubated 1 h and the monomer was isolated by gel filtration in 20 mM sodium phosphate buffer, pH 8.0. Two molar equivalents of Alexa-647 C_2_ maleimide (Thermo Fisher A20347) were added from a concentrated stock in DMSO. The solution was incubated overnight in darkness on ice. Labelled monomer was isolated from free dye by two rounds of gel filtration in 20 mM sodium phosphate buffer, pH 8.0. Antibodies were labelled by mixing with Alexa-647 N-hydroxy succinimidyl ester (Thermo Fisher A20006) after gel filtration of each antibody in PBS. Two molar equivalents of Alexa-647 were added to each antibody, and the solutions were incubated for 2 h at 4°C, followed by gel filtration twice to remove free dye. The absence of free dye was confirmed using microfluidic diffusional sizing with fluorescence detection using a Fluidity One-W instrument (Fluidic Analytics Ltd, Cambridge, UK)

### 1.14 Binding measurements using microfluidic diffusional sizing (MDS)

Binding interactions in solution were assessed through diffusion measurements under laminar flow in a microfluidic device using a Fluidity One-W instrument (Fluidic Analytics Ltd, Cambridge, UK). The smaller of the two interacting species was used in labelled format. Thus Alexa-647-Aβ42 was used to study interactions between peptide monomers and antibodies, and Alexa-647-antibodies were used to study the interaction between antibodies and Aβ1-42 fibrils. Alexa-647-Aβ(MC1-42) monomer was isolated by SEC and flash-frozen or kept on ice until use. Unlabelled Aβ1-42 fibrils were formed quiescent at 37 °C in PEGylated plate wells (Corning 3881) from monomer. After formation the fibrils were sonicated (20/20 sec break cycles) for 6 min on ice and shaken for 30 min at 1800 rpm to provide ca. 50 nm diffusible fibrils for the binding experiments. The labelled species was combined with unlabelled binding partner, and incubated for 15 minutes before the diffusion measurements. All binding measurements were performed at 27-28 °C in PBS, pH 7.8. The sample was injected in one half of the channel and at the outlet, the fractional fluorescence intensity in the two halves of the channel were analysed separately, *f*_*N*_ representing the fraction of fluorescent species in the non-diffused half and *f*_*D*_ the fraction of fluorescent species in the diffused half. Each condition was repeated 3-7 times and the data are shown in the form of all individual replicates.

### 1.15 Fitting to microfluidics diffusion data

Equilibrium binding parameters (affinity and stoichiometry) were obtained by fitting directly to the raw data in terms of the fraction of fluorescent species appearing in the diffused half of the channel. Because relatively dilute solutions were used, activities were replaced with concentrations and the apparent dissociation constant, *K*_*D*,_ was estimated by fitting to the measured fraction of fluorescent species appearing in the diffused half of the channel, *y*, using the following equation for *f*_*D*_, the calculated fraction of fluorescent species appearing in the diffused half of the channel

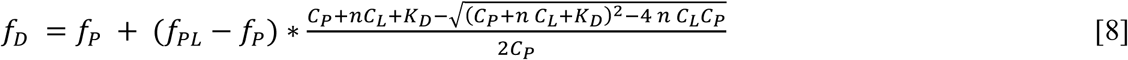

where *C*_*P*_ is the total concentration of labelled protein (monomer or antibody), *C*_*L*_ is the total concentration of unlabelled protein (the ligand), *f*_*P*_ and *f*_*PL*_ are the fractions of free labelled protein and bound labelled protein, respectively, in the diffused half of the channel, and *n* is the stoichiometry of the interaction, such that *nC*_*L*_ is the total concentration of binding sites on the unlabelled component. For data with Alexa-647-labelled Aβ42 monomer, *n* was fixed to 2 (two monomer-binding sites per antibody), while *f*_*P*_, *f*_*PL*_ and *K*_*D*_ were fitted parameters. For data with Alexa-647-labelled antibodies, *n, f*_*P*_, *f*_*PL*_ and *K*_*D*_ were fitted parameters.

Based on the ratio of *f*_*N*_ and *f*_*D*_, the instrument reports an apparent hydrodynamic radius of the fluorescently labelled species, the principles of which have been described previously (*20, 21*). However, the measured radius is not linearly related to the concentrations of free and bound labelled substrate.

#### Bayesian inference details

The Bayesian inference analysis method utilizes Bayes’ theorem, and allows the determination of the probability distributions of unknown parameters, given the observed data, by the following equation:

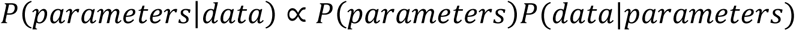

where *P*(*parameters*|*data*) is known as the posterior, *P(parameters)* as the prior, and *P*(*data*|*parameters*) as the likelihood. The prior probability distribution is an expression of our information about the system before we acquire any measurement data. For *f*_*P*_ and *f*_*PL*_, we assume the prior is flat in linear space, whereas for the *K*_*D*_ and, in the case of the fibrils, *n*, a prior that is flat in logarithmic space is more appropriate, to reflect the scale invariance of the problem.

We assume our experimental measurement data to be normally distributed about the true value; our likelihood function is therefore a Gaussian, centered on the theoretical measurement value.

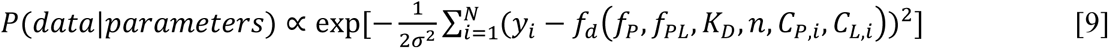

where *f*_*d*_ is defined as above, with or without the n dependence, as determined by whether the Aβ is monomeric or fibrillar, *C*_*P,i*_ and *C*_*L,i*_ are the total concentrations labelled and unlabeled species, respectively, in the i^th^ measurement, with both having an index i as we combine all data sets for each combination of labeled and unlabeled species. *y*_*i*_ is the fraction of diffused labelled component in the i^th^ measurement. In order to define an appropriate standard deviation, *σ*, for each dataset, we calculate the standard deviations of repeats of each measurement, and use the maximum of these values as a global standard deviation for that dataset.

### 1.16 Fraction bound labeled species

After fitting, the fractional saturation of the labelled species, *Q*, was calculated from each data point, using the fitted values of *f*_*P*_ and *f*_*PL*_ as follows

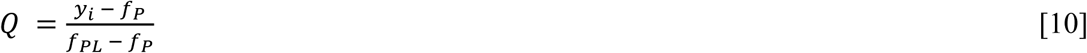

and plotted versus free ligand concentration, which was calculated at each total concentration of added ligand, *C*_*L*_, as follows

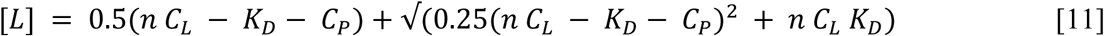

using in the case of Alexa-647-Aβ42 interacting with antibody the fitted value of *K*_*D*_ and the fixed values of *n* and *C*_*P*_, and in the case of Alexa-647-antibody interaction with fibrils, the fitted values of *n* and *K*_*D*_ and the fixed value of *C*_*P*_.

### 1.17 Fraction of fibrils with at least one antibody bound

The fraction of fibrils with at least one antibody bound, f_B_, was calculated assuming no cooperativity, i.e. the affinity for each site is the same irrespective of whether the other sites on the same fibril are occupied or not. The probability of a given site to be occupied is 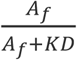, where *A*_*f*_ is the concentration of free antibody. The number of species bound per fibril gives a binomial distribution, thus the fraction of fibrils with one or more antibodies bound is given by

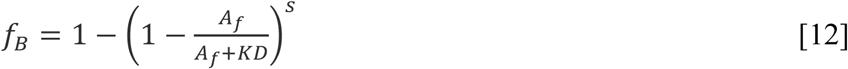

where *s* is the number of antibody binding sites on the fibril and we used the fact that the fraction of fibrils that have more than one antibody bound is equivalent to 1 minus the fraction that has no antibodies bound. The values used to calculate the curves shown in Figure 4L assume short fibrils of 400 monomers (50 nm fibrils, two filaments, two monomers per plane in each filament (22):

| Antibody | Monomers /<br>antibody | $s$ | $K_D$ / nM |
| --- | --- | --- | --- |
| <sup>ch</sup>aducanumab | 5 | 80 | 1 |
| 3D6 | 47 | 8 | 0.27 |
| <sup>ch</sup>gantenerumab | 43 | 9 | 30 |

